# Investigating the effect of ginger-derived nanovesicles on the growth and metabolic activity of *Bacteroides thetaiotaomicron*: an isothermal microcalorimetric study

**DOI:** 10.64898/2026.01.08.698378

**Authors:** Francesca Loria, Anna Kattel, Marco Brucale, Francesco Valle, Paolo Bergese, Eeva-Gerda Kobrin, Antonio Chiesi, Paolo Guazzi, Sirin Korulu, Irina Stulova, Nataša Zarovni, Raivo Vilu

## Abstract

Plant-derived extracellular vesicles (EVs) have shown numerous health benefits, including modulation of the human gut microbiota. Hereby, we employed isothermal microcalorimetry (IMC) to explore the effects of ginger-derived nanovesicles (G-NVs) on the growth and metabolic activity of *Bacteroides thetaiotaomicron (Bt),* a dominant saccharolytic commensal with promising next-generation probiotic potential. *Bt* was exposed to either G-NVs or the ginger extract separated from G-NVs (G-CTL) in defined media under anaerobic conditions. Both ginger samples enhanced bacterial specific growth rate and maximum metabolic activity, inducing the latter earlier. However, higher biomass production and greater secretion of acetic, succinic, and propionic acids occurred only in response to the G-CTL. Complete sugar depletion and unchanged FAA levels indicated preferential carbohydrate utilization by *Bt*. Overall, these findings revealed that *Bt*’s metabolic state is shaped by both G-NVs and G-CTL, yet through distinct mechanisms, with G-NVs inducing rapid stimulation without increasing total metabolic output and G-CTL providing a sustained, dose-dependent effect. To our knowledge, this is the first study applying IMC to monitor real-time the impact of EVs on microbial growth and metabolism, underscoring IMC’s utility for mechanistic studies of EV-microbe interactions. Furthermore, this research sets the ground for innovative strategies in nutraceutical and microbiome-targeted therapy development.

## 1. Introduction

The human gastrointestinal tract hosts a diverse collection of coexisting microorganisms, including bacteria, archaea, and viruses, whose composition and functionality critically regulate human health [1]. The microbial communities residing in the intestine, collectively termed human gut microbiota (HGM), are considered key regulators of host homeostasis due to their high microbial load, metabolic activity, immune functions, and systemic impact [2,3]. Indeed, in addition to supporting nutrient digestion, metabolism, and gut barrier integrity, HGM is shown to play an important role in promoting direct regulation of the host immune cells and fostering protection against the action of pathogens and toxins [4–7]. Recent studies have also provided evidence of its prominent action in modulating the development and proper functioning of the nervous system via the gut-brain axis. The HGM is mainly composed by the following phyla: Bacteroidetes, Firmicutes, Actinobacteria, Proteobacteria, and Verrucomicrobia [8]. In adult individuals, the Gram-negative Bacteroidetes and the Gram-positive Firmicutes constitute more than 80% of the entire microbial community [9]. Within the Bacteroidetes phylum, the genus *Bacteroides* has been extensively studied due to its dominance over other genera and its long-term adaptability to the changing host’s gut environment across human populations [10].

*Bacteroides thetaiotaomicron (Bt)* is a non-motile, obligate anaerobe primarily found in the large intestine. By virtue of its prevalence, functional relevance, metabolic flexibility, and genetic tractability, *Bt* is considered an ideal model organism for studying the HGM. Its biological significance mainly relies on its direct regulation of the host gene expression pertaining to metabolism, immunomodulation, and maintenance of the gut barrier integrity [11]. *Bt* shares versatile metabolic capabilities characteristic for all *Bacteroides* spp. Upon carbohydrate fermentation, *Bt* secretes several organic and inorganic products that support its primary biological functions, including short-chain fatty acids (SCFAs) (*e.g.*, acetate, propionate, and formate), succinate, ethanol, lactic acid, CO_2_, and H_2_. As *Bt* is capable of fermenting proteins and amino acids under specific conditions, it has also been demonstrated that *Bt* releases SCFAs, along with branched-chain fatty acids, phenolic/indolic compounds, amines, and ammonia [12]. Technically, *Bt* is easy to cultivate under anaerobic conditions and serves as an excellent candidate for HGM-focused co-cultured studies, as it interacts with other commensal species in a predictable manner [13–16]. With its whole genome sequenced, *Bt* can also be genetically manipulated, thus contributing to facilitate fast-paced advancements in the synthetic biology and microbiome (engineering) fields [17]. Because of these important attributes, *Bt* shows significant potential as next-generation probiotic (NGP) [18,19]. Probiotics are viable and beneficial microorganisms exerting health-enhancing and health-protecting properties when administered in sufficient quantities. Compared with traditional probiotics, including *Lactobacillus* and *Bifidobacterium* spp., NGPs demonstrate superior metabolic benefits and offer the competitive advantage of targeting specific disease conditions, rather than managing overall health [20].

Biological nanoparticles, classified as extracellular vesicles (EVs), have recently been documented in plants as bioactive components exhibiting health-protecting and enhancing properties (*e.g.*, anti-cancer, anti-inflammatory, antioxidant, and prebiotic) [21–23]. Plant-derived EVs and nanovesicles (*i.e.*, these last indicating co-isolates of both natural EVs and artificial particles resulting from mechanical and destructive pre-processing of the plant source [24]) are, hereby, collectively referred to as plant-derived vesicles (PDVs). PDVs are lipid-bilayer membranous particles with nanometer and micrometer size: ^∼^50-150 nm, defined as “small EVs"; ^∼^200-800 nm, defined as "medium EVs", and ≥ 1,000 nm, defined as "large EVs" [23–25]. By carrying diverse bioactive components (*e.g.*, proteins, lipids, nucleic acids, and metabolites), PDVs were shown to mediate intercellular communication within and across organisms. Additionally, they have demonstrated to be involved in plant cell proliferation and differentiation, cell wall remodeling, and defense mechanism against abiotic and biotic stresses. Due to their inner composition and functional properties, PDVs are rapidly gathering momentum as competitive clinical-grade and commercial-grade tools that can be deployed between several industrial markets (*e.g.*, healthcare, personal care, and agrifood) [26]. Among the PDVs that have thus far promoted important health benefits in preclinical settings, small ginger rhizome (*Zingiber officinale Roscoe*)-derived nanovesicles (G-NVs) have demonstrated significant potential in regulating and fostering a healthy HGM [27,28].

Isothermal microcalorimetry (IMC) enables real-time monitoring of heat changes associated with complex biological processes in closed systems (*e.g.*, ampoules) of small caliber at a constant temperature [28]. For decades, IMC has been widely applied to study bacterial growth and metabolism, providing quantitative and thermodynamic insights into growth dynamics, kinetics, and metabolic activity. This technique measures the heat generated by microbial metabolic activity and quantitatively relates it to biomass formation, substrate consumption, and carbon source utilization efficiency [30–35]. Compared with traditional methods, IMC offers high robustness, throughput, sensitivity (*i.e.*, a limit of detection ranging from 10^4^ to 10^5^ cells), and reproducibility. Furthermore, it does not require optical transparency, specific physical forms, (pre-)treatment, or any special preparation of the sample(s) prior to assessment [36–38]. Despite these advantages, IMC has not yet been reported as a method for investigating the influence of EVs on microbial growth.

The primary aim of this study was to apply IMC to explore the effect of G-NVs on the growth and metabolic activity of the human gut bacterium *Bt*. This was analyzed by measuring the dose-dependent functional activity of G-NVs, enriched in the sterile retentate of a tangential flow filtration (TFF) system with 50 nm pores. The ginger extract depleted of the G-NV sample, equivalent to the TFF permeate enriched in constituents smaller than 50 nm and mostly non-vesicular, was assessed in parallel as a procedural control sample (G-CTL). The metabolic response of *Bt* to the treatment with G-NVs and G-CTL was further evaluated by high-performance liquid chromatography (HPLC) and Ultra Performance Liquid Chromatography (UPLC) with Ultraviolet/Visible or Refractive index detection analysis of non-vesicular extracellular metabolites [*i.e.*, sugars, organic acids, ethanol, and free amino acids (FAAs)] secreted throughout microbial growth (Figure 1). Our data provided evidence of the great potential of IMC to unravel the dynamic metabolic regulation of *Bt* upon exposure to ginger sample fractions, specifically showing that both G-NVs and G-CTL affected *Bt*’s growth and metabolism, yet through different mechanisms and kinetics.

**Figure 1.**
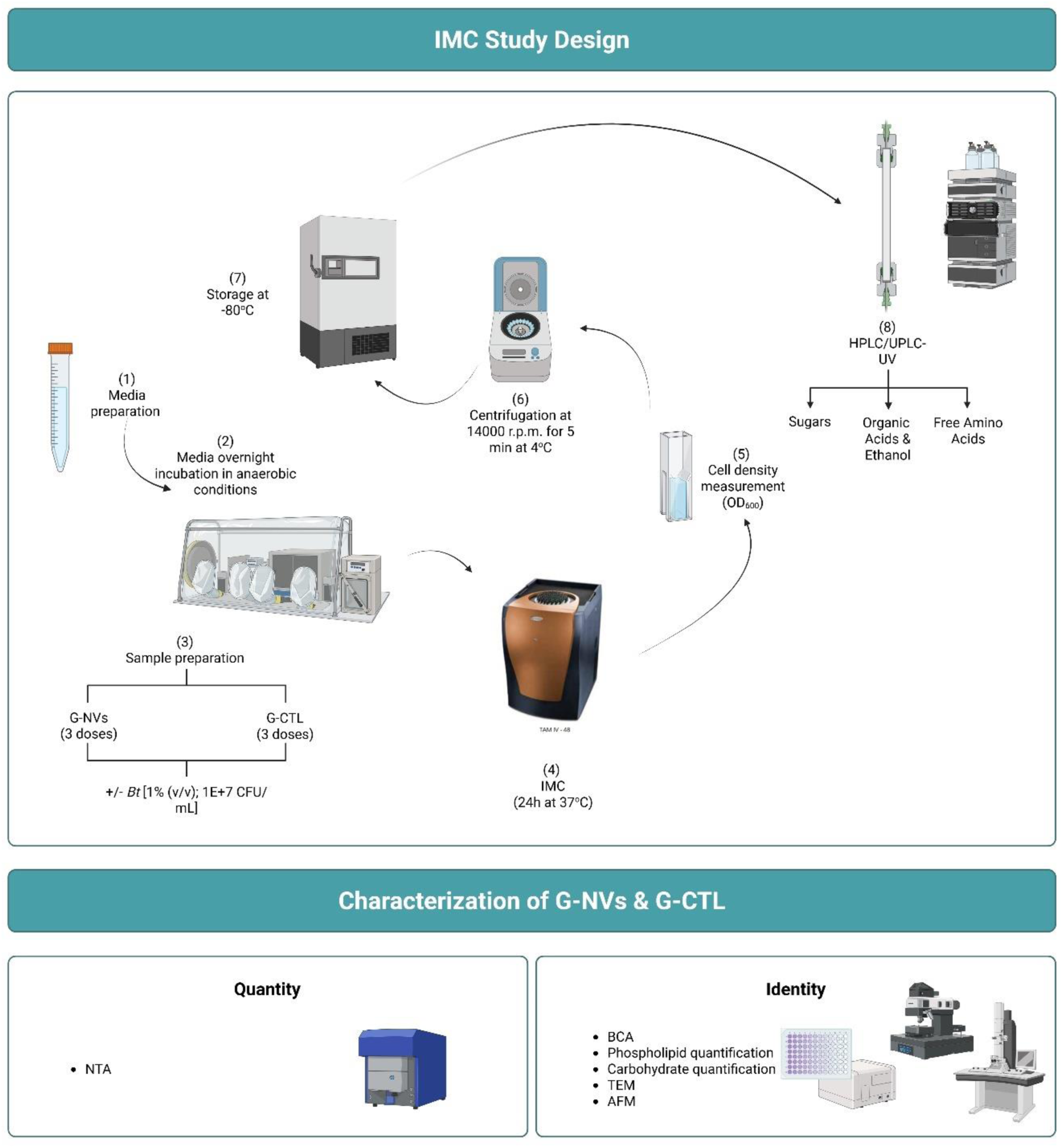
Graphical overview of the study design. Legend: AFM = atomic force microscopy; CFU = colony-forming units; G-CTL = G-NVs procedural control; G-NVs = ginger-derived nanovesicles; HPLC = high-performance liquid chromatography; IMC = isothermal microcalorimetry; OD_600_ = optical density at a wavelength of 600 nm; TEM = transmission electron microscopy; UPLC-UV = ultra-performance liquid chromatography-UV. This figure was created with BioRender.com.

## 2. Materials and Methods

### 2.1. Production of G-NVs and G-CTL

Ginger rhizome (*Zingiber officinale Roscoe*) of Chinese origin was obtained from a local Estonian market. Ginger rhizome was washed with Milli-Q water, followed by peeling, slicing, and grinding with a centrifugal juicer (Severin; #ES3566) to extract juice and dry residues. For each batch, 30 g of dry residues were resuspended in 60 mL of pre-extracted juice supplemented with protease inhibitor cocktail (SigmaFast^TM^ Protease Inhibitor Tablets; Sigma-Aldrich; #S8820; 50X working dilution) and diluted with phosphate buffered saline (PBS) (Corning; #21-031-CV) up to 300 mL. After 30 minutes of gentle shaking at room temperature (RT), residues were discarded using a funnel and a drainer. Then, samples were centrifuged twice at RT, using a Ohaus FC5718R centrifuge with fixed rotor: i) at 300 r.c.f. for 20 min and ii) at 3,000 r.c.f. for 20 min. The resulting supernatant was subsequently subjected to vacuum-based glass filtration with a glass fiber prefilter (Merck; #APFD09050). After dead-end filtration through a 0.45 µm pore size filter and a further centrifugation step at 10,000 r.c.f. for 30 min at RT, samples were processed by TFF, using TFF-EVs-Small device (HansaBioMed Life Sciences; #HBM-TFF-EVs-S). Recovered TFF retentate and permeate were finally sterilized by dead-end filtration using 0.22 µm pore size filter units (Sartorius; #16532). Finally, the filtered TFF retentate and permeate were stored at −80°C to be subsequently analyzed as small ginger-derived nanovesicles (G-NVs) and G-NVs procedural control (G-CTL), respectively.

### 2.2. Characterization of G-NVs and G-CTL

#### 2.2.1. Nanoparticle Tracking Analysis (NTA)

Nanoparticle concentrations and size distributions were analyzed by Nanoparticle Tracking Analysis (NTA), using Particle Metrix ZetaView PMX-120 (software version 8.05.12 SP1), as previously described [39].

#### 2.2.2. Fluorescence Labeling

Fluorescence labeling was optimized and conducted using CellMask™ Green (CMG) Plasma Membrane Stain (Thermo Fisher Scientific; #C37608) [peak excitation wavelength (λex) = 522 nm; peak emission wavelength (λem) = 535 nm], as shown prior [39]. Briefly, for each G-NV biological batch, 5E+9 particles – quantified by NTA in scatter mode – were incubated with CMG (1:50 dilution or 20X working concentration), in a final reaction volume of 25 µL, for 90 minutes at 37°C in the dark. The G-CTL was stained at the same dilution as its corresponding G-NV sample. After staining, the excess of unbound dye was removed by size-exclusion chromatography (SEC) using miniPURE-EVs SEC columns (HansaBioMed Life Sciences; #HBM-mPEV-##), according to the manufacturer’s datasheet. The eluate corresponding to the EV fraction (500 µL) was then assessed by fluorescence NTA (F-NTA). Three replicates of a fluorescence labeling control sample, equivalent to the buffer-only (PBS) supplemented with 20X CMG, was analogously processed and analyzed in parallel to assess the effectiveness of SEC in removing the excess of unbound dye.

#### 2.2.3. Fluorescence Nanoparticle Tracking Analysis (F-NTA)

F-NTA was conducted using Particle Metrix ZetaView**®** Evolution (software version 1.0) equipped with a CMOS-camera sensor and lasers of 405 nm (violet), 488 nm (blue), 520 nm (yellow/orange), and 640 nm (red) wavelengths. Instrument setup was conducted according to the manufacturer’s instructions. Briefly, alignment of the optical system (camera and lasers), focus optimization, and temperature equilibration (22–25°C) were performed using 100 nm polystyrene beads (Applied Microspheres; #75009-03) and fluorescent PS OR-520 beads for 520 nm laser (Particle Metrix; #700093). Prior to measurement, CMG-labelled samples and procedural controls were diluted 1:10 and 1:20 in PBS to achieve an optimal scattering particle count of 50-200 particles per cell position throughout a total of 11 cell positions. Scatter measurements were performed in the scatter channel illuminated with a 488 nm (blue) laser (SC488), selecting a camera gain of 40 and a shutter speed of 200. These operational settings were selected to align readouts with those previously obtained using Particle Metrix ZetaView PMX-120. Fluorescence measurements were conducted in the fluorescence channel illuminated with a 520 nm (yellow/green) laser (FL520) and coupled with a 550 nm cut-on long-pass fluorescence filter, adjusting the camera gain to 80 and maintaining the shutter speed 200. Each size and concentration measurement, both in scatter and fluorescence mode, was performed in triplicate. Data were presented as mean ± standard error of the mean (SEM) from the three technical measurements.

#### 2.2.4. Bicinchoninic Acid (BCA) assay

Proteins were quantified by Bicinchoninic Acid (BCA) assay, using the Pierce™ BCA Protein Assay kit (Thermo Fisher Scientific; #23228), according to the manufacturer’s instructions. Standards and samples were loaded in duplicate into transparent Nunc™ 96-Well Polystyrene Round Bottom Microwell Plates (Thermo Scientific; #268152). Optical densities (ODs) were measured at 540 nm using Tecan GENios Pro microplate reader.

#### 2.2.5. Phospholipid assay

Phospholipid concentrations were measured using fluorescent Phospholipid Quantification Assay Kit (HansaBioMed Life Sciences), following the manufacturer’s protocol. Standards, samples, blanks, and background controls were assessed in duplicate into black polystyrene - no binding capacity – microplate wells (HansaBioMed Life Sciences). Fluorescence intensities (FIs) were read using Tecan GENios Pro microplate reader (λex = 540 nm; λem = 590 nm).

#### 2.2.6. Total carbohydrate assay

Total carbohydrate quantification was performed by Total Carbohydrate Colorimetric Assay Kit (Sigma-Aldrich; #MAK104-1KT), according to the manufacturer’s datasheet. Standards, samples, and blanks were analyzed in duplicate, using transparent polystyrene - no binding capacity – microplate wells (HansaBioMed Life Sciences). ODs were read at 492 nm, using Tecan GENios Pro microplate reader.

#### 2.2.7. Transmission Electron Microscopy (TEM)

For morphological assessment, transmission electron microscopy (TEM) was performed as done prior [40]. Briefly, samples were prepared by loading to carbon coated and glow discharged 200 mesh copper grids with pioloform support membrane. Then, samples were fixed with 2.0% PFA in NaPO_4_ buffer, stained with 2% neutral uranyl acetate, further stained, and embedded in uranyl acetate and methyl cellulose mixture (1.8/0.4%). Finally, samples were visualized using TEM Hitachi HT7800 operating at 100 kV. Images were taken with Gatan Rio9 bottom mounted CMOS-camera, model 1809 (Gatan Inc., USA) with 3072 x 3072 pixels image size and no binning.

#### 2.2.8. Atomic Force Microscopy (AFM)

Atomic force microscopy (AFM) and subsequent quantitative morphometry were performed on vesicular samples as previously described [41,42]. Briefly, images were taken in PeakForce mode on a Multimode 8 microscope (Bruker, USA) equipped with Scanasyst Fluid+ probes (Bruker), a Nanoscope V controller, a JV-type piezoelectric scanner, and a sealed fluid cell. For each sample, 5 μL aliquots were loaded on poly-L-lysine-coated glass coverslips for 30 minutes at 4°C to promote adsorption, followed by direct introduction into the fluid cell without prior drying. To maximize the number of isolated particles, thus facilitating automatic image analysis, sample concentration was adjusted through successive depositions. For proper comparative assessment, G-NV and G-CTL samples, at the same dilution, were deposited, scanned, and analyzed following an identical pipeline. Quantitative morphometric analysis was performed using Gwyddion 2.61 [43] and custom Python scripts. Each particle on the surface was assessed for its spherical diameter in solution (D), as well as for its surface contact angle (CA), which has previously demonstrated to directly correlate with the mechanical stiffness of intact EVs. For each sample, all measured particles were plotted on CA vs. D scatterplots. Intact vesicles and non-vesicular particles were previously shown to cluster into characteristic zones of these plots [41, 42].

### 2.3. Assessment of bacterial growth and metabolic activity

#### 2.3.1. Media preparation

The experiment was conducted using a pre-defined mineral medium, whose composition consisted of: D-(+)-Glucose monohydrate (2.2 g/L), L-amino acids [(g/L) Ala 0.11, Arg 0.0575, Asn 0.095, Asp 0.095, Glu 0.09, Gln 0.045, Gly 0.08, His 0.0675, Ile 0.15, Leu 0.3, Lys-HCl 0.2, Met 0.0575, Phe 0.125, Pro 0.1025, Ser 0.2375, Thr 0.1025, Trp 0.0225, Val 0.15, Tyr 0.0375], mineral salts [(mg/L) MgSO_4_ × 7H_2_O 36, MgCl_2_ 2, FeSO_4_ × 7H_2_O 0.1, CaCl_2_ 9, MnSO_4_ × H_2_O 3, ZnSO_4_ × 7H_2_O 1, CoSO_4_ × 7H_2_O 1, CuSO_4_ × 5H_2_O 1, (NH_4_)6Mo_7_O_24_ × 4H_2_O 1, NaCl 527], (NH_4_)Cl 3.5; NaHCO_3_ 1], vitamins [(mg/L) biotin 6.02, riboflavin 0.9, pyridoxamine-2HCl 5, niacin 0.9, thiamine-HCl 0.56, pyridoxine-HCl 4.8, Ca-pantothenate 1.2, folic acid 0.56, p-aminobenzoic acid 0.05, lipoic acid 1.05, ascorbic acid 500, vitamin B12 0.001, vitamin K (menadione) 1], hemin (5 mg/L). Media were prepared using 1M dipotassium hydrogen phosphate buffer (16.8 mL/L) and 1M potassium dihydrogen phosphate buffer (34.2 mL/L), and supplemented with L-Cysteine hydrochloride (1 g/L) as a sulfur source and reducing agent. Media were filtered with filter units having 0.22 µm pore size, followed by overnight incubation in an anaerobic chamber (COY box, Coy Laboratory Products Inc.) with an atmosphere of 2.5 ± 0.5% H_2_, 10% CO_2_, and balanced N_2_. If needed, the pH of the media was adjusted to 7.0 ± 0.1 before performing the experiment.

#### 2.3.2. Assessment of bacterial growth activity by isothermal microcalorimetry (IMC) and optical density measurement

The *Bt* stock culture was centrifuged at 19,000 r.c.f. for 3 min at RT, followed by pellet resuspension in PBS solution containing 1% L-Cysteine hydrochloride. Cell counting was adjusted to a working stock concentration of 1-1.2E+9 colony-forming units (CFU)/mL. For each sample supplemented with or without G-NVs and G-CTL, 10 or 15 µL of inoculum [1% (v/v); final cell concentration of 1-1.2E+7 CFU/mL was added to 990 or 1485 µL of medium, respectively. Treated bacteria were exposed to three distinct doses of G-NVs (*i.e.*, 2E+11, 1E+11, and 5E+10 scattering particles) and G-CTL, which was equivalent to the corresponding G-NV dose based on protein content. To exclude microbial contamination, equal doses of non-inoculated G-NVs and G-CTL were tested in parallel. These contained 990 or 1485 µL of medium with either G-NVs or G-CTL, which was supplemented with 10 or 15 µL of PBS [1% (v/v)], respectively. All samples were run in triplicate in four independent experiments. Each experiment assessed a distinct biological batch of both G-NVs and G-CTL. After being hermetically sealed, the vials were placed into a TAM IV-48 (48-channels, TA Instruments) heat conduction multi-channel microcalorimeter to assess bacterial growth and growth kinetics. The experiment was run at 37°C up to 24 hours. Subsequently, the samples were transferred into new Eppendorf tubes, and the cell density was measured as OD at a wavelength 600 nm (OD_600_) using an Ultrospec 10 Cell Density Meter (Biochrom). As described elsewhere [44], the biomass concentration (gDW L^-1^) was estimated using OD_600,_ according to the following formula:

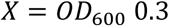

Finally, after centrifugation of the samples at 19,000 r.c.f. for 5 min at 4°C to separate the cells from conditioned media, the resulting supernatants were transferred to new Eppendorf tubes. Both cell pellets and supernatants were stored at −80°C for subsequent analyses.

##### 2.3.2.1. Analysis of IMC power-time curves and calculation of growth characteristics

A typical power-time (or heat flow) curve relative to the growth of individual microbial species, such as *Bt*, describes the evolution of heat released by bacteria throughout the entire growth process. Specifically, it illustrates a thermogram plotting the rate of heat against the unit of growth time (W). As in any microbial growth curve, which plots the biomass concentration against the unit of growth time, primary, consecutive growth phases can be discriminated: i) lag phase (*i.e.,* no rise in number of living bacterial cells due to initial adjustment to the new environment); ii) exponential phase (*i.e.,* exponential rise in number of living bacterial cells); iii) deceleration phase (*i.e.*, plateau in number of living bacterial cells due to equal rate of bacterial cell division and death, followed by metabolic slowdown).

The bacterial metabolic evolution displayed by the power-time curve may be described through thermodynamic and kinetic parameters that can be analytically determined through mathematical analysis of the curve: i) the maximum specific growth rate (*µ_max_*), which is the rate at which a microbial population increases in size per unit of time; ii) the maximum power output or metabolic activity level (*P_max_*) at the highest peak of the calorimetric curve; iii) the time at which the *P_max_* occurs (*T_Pmax_*); iv) the cumulative heat produced during the exponential phase (*Q_exp_*); v) the cumulative heat produced during the whole cultivation period (*Q_tot_*).

The *µ_max_* is determined based on the direct relationship between the biomass concentration (*X*), the rate of biomass concentration 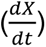, and the specific growth rate (*µ),* satisfying the following mathematical equation [45]:

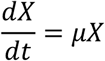

During the exponential growth phase, the biomass increases exponentially with time, as described below:

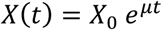

in which *X(t)* is the biomass at time point *t* of the exponential growth phase and *X_0_* is the biomass at time point 0 of the exponential growth phases.

Based on the assumption that the rate of biomass production is directly proportional to the rate of heat production 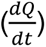, the *µ_max_* can be derived from the slope of the exponential phase of the relative power-point curve, as follows:

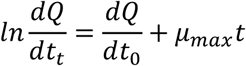

in which 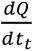 is the rate of heat production at the time point *t* of the exponential growth phase and 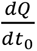 is the rate of heat production at a time point 0 of the exponential growth phases.

The *Q_exp_* and *Q_tot_* are mathematically expressed as the integrals of the power-time curve over specific growth intervals: i) between the initial time point and the final time point of the exponential growth phase (*Q_exp_*); ii) between the initial time point and the final time point of the entire course of the microbial growth (*Q_tot_*). *Q_exp_* is depicted by the area under the power-time curve comprised between the baseline and an imaginary vertical line passing through the peak of the power-time curve (*P_max_*) and intersecting the x-axis at time point *T_Pmax._* In contrast, *Q_tot_* corresponds to the area comprised between the power-time curve and the baseline over the total duration of the microbial cultivation [32–35].

Raw data were collected from the TAM Assistant program (version 0.9.1012.40, SciTech Software AB, Thermometric AB, Järfälla, Sweden). Heat evolution was recorded at a frequency of 1 s^−1^ and then resampled at 1-hour intervals. All parameters were estimated through Microsoft Excel [32–35]. Baseline-corrected integrals of the power-time curve, which were automatically provided by the software, were used to calculate *Q_exp_* and *Q_tot_*.

#### 2.3.3. Analysis of bacterial extracellular metabolites

##### 2.3.3.1. Liquid Chromatography (LC)

Fermentation supernatants were centrifuged at 21,100 r.c.f. for 6 min at 4°C. The supernatant was transferred into the prerinsed with Milli-Q Amicon Ultra 0.5ml 3 kDa Molecular Weight Cut-off spin filters (Merck Millipore Ltd; #UFC5003) and centrifuged at 6,150 r.c.f. for 30 min at RT. Filtered samples were 10 times diluted with Milli-Q water and assessed by high-performance liquid chromatography (HPLC) to determine the content of sugars [i.e., glucose (Glc), fructose (Fru), mannose (Man), N-acetyl-D-glucosamine (GlcNAc), rhamnose (Rha)], organic acids (*i.e.*, acetate, butyrate, propionate, malate, formate, fumarate, glucuronic acid, isobutyrate, isovalerate, lactate, succinate), and ethanol, using Aminex HPX-87H columns (Bio-Rad Laboratories; #1250140). Ultra Performance Liquid Chromatography-UV (UPLC-UV) was adopted to examine and quantify FAAs (*i.e.*, alanine (Ala), arginine, (Arg), asparagine (Asn), aspartic acid (Asp), cysteine (Cys), glutamic acid (Glu), glycine (Gly), histidine (His), isoleucine (Ile), leucine (Leu), lysine (Lys), methionine (Met), phenylalanine (Phe), proline (Pro), serine (Ser), threonine (Thr), tryptophan (Trp), tyrosine (Tyr), valine (Val). Ammonium ion (NH_4_+) levels were also analyzed in parallel, as being a key indicator of amino acid metabolism. Both HPLC and UPLC-UV analyses were conducted as previously described [44].

### 2.4. Statistical analyses

Statistical analysis was conducted using GraphPad Prism 10 (GraphPad Software, San Diego, CA, USA). Data groups were compared by either ordinary unpaired T test, one-way ANOVA, or two-way ANOVA with follow-up test for multiple comparisons, as outlined in the descriptive captions of each figure. Statistical significance was claimed when the p-values were < 0.05 [*], <0.01 [**], <0.001 [***], and <0.0001 [****]. For each sample, data were presented as mean ± SEM from three technical replicates, calculated across three to four independent biological batches, as described in the descriptive captions of each figure.

## 3. Results

### 3.1. Physicochemical assessment of G-NVs and G-CTL

Prior to the IMC study, we assessed G-NVs and G-CTLs for their physicochemical properties. NTA revealed that the G-NV preparation was enriched in particles with median and peak sizes of 97.5 ± 4.4 and 98.9 ± 6.0 nm, respectively. Particles have been also detected in the G-CTL, yet at significantly lower concentration and revealing a size distribution shifting towards smaller sizes (Figures 2A-B and S1A-B).

**Figure 2:**
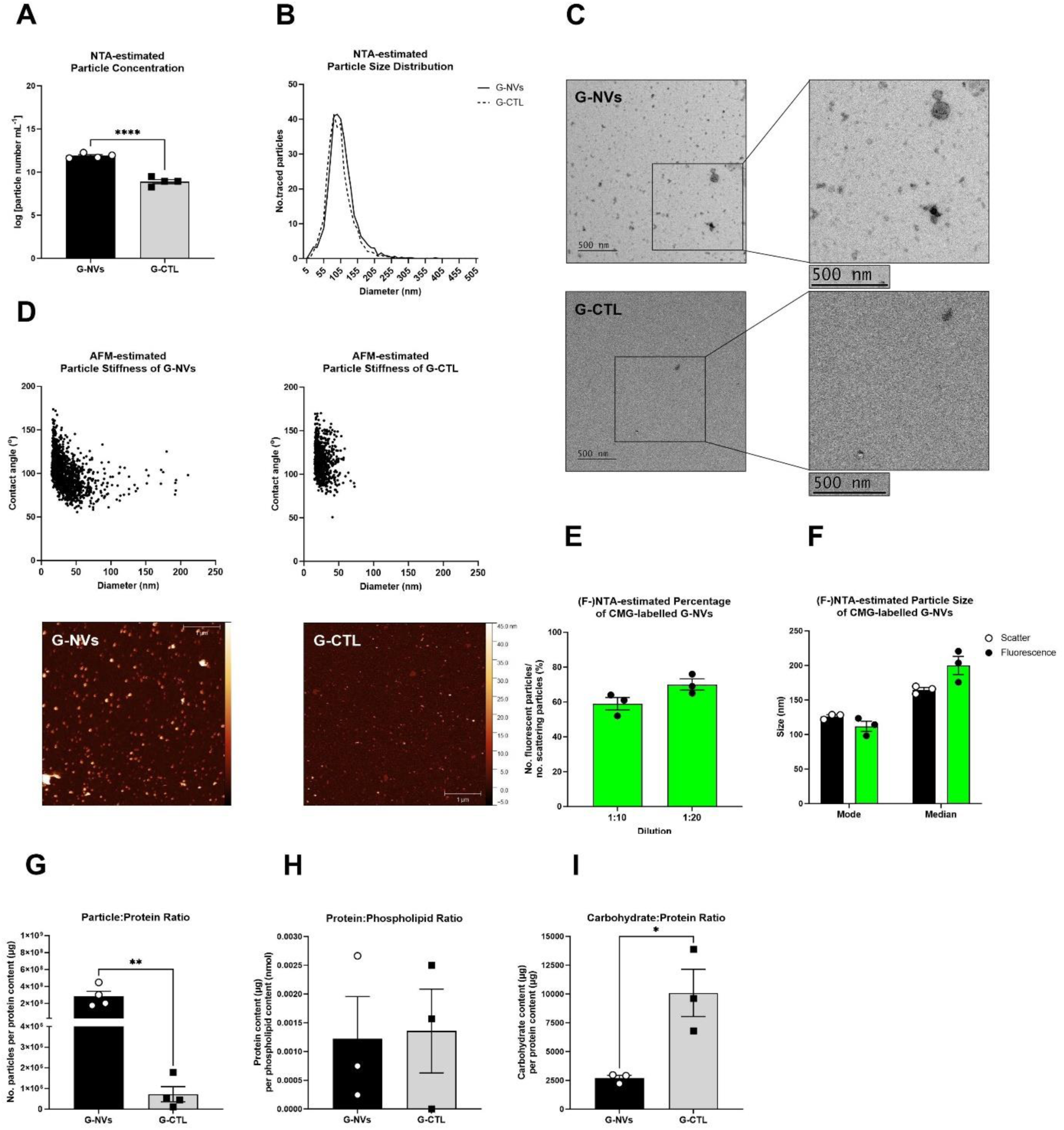
Physicochemical assessment of G-NVs and G-CTL. The bar graphs show: A) NTA-estimated particle concentrations of four biological batches of G-NVs and G-CTL – expressed as particle number per milliliter in logarithmic scale; E) (F-)NTA-estimated ratio of total number of fluorescent particles over total number of scattering particles detected in three biological batches of CMG-labelled G-NVs – expressed as percentage (%) and measured with two serial dilutions (1:10 and 1:20) in PBS; F) (F-)NTA-estimated particle mode size (nm) and median size (nm) of three biological batches of CMG-labelled G-NVs measured in scattering and fluorescence mode with 1:10 dilution in PBS; G) particle:protein ratio of four biological batches of G-NVs and G-CTL – expressed as particle number per protein content (µg); H) protein:phospholipid ratio of three biological batches of G-NVs and G-CTL – expressed as protein content (µg) per phospholipid content (nmol); I) carbohydrate:protein ratio of three biological batches of G-NVs and G-CTL – expressed as carbohydrate content (µg) per protein content (µg). The histograms in B depict the NTA-estimated particle size distributions of four biological batches of G-NVs and G-CTL – expressed as average of two to four technical measurements per sample. The (C) TEM and (D) liquid AFM images illustrate, respectively, the morphological and nanomechanical properties of one representative batch of G-NVs and G-CTL. The scatterplots in D show the contact angle (°) and equivalent diameter (nm) of each particle detected and analyzed by liquid AFM. Bargraph data are reported as mean ± SEM. In A, E, G, H, and I, statistical analysis was performed using ordinary unpaired T test (not reported p-value: not significant; significant p-value: < 0.05 [*], <0.0001 [****]). In F, statistical analysis was performed between matching scatter and fluorescence values using using ordinary unpaired T test (not reported p-value: not significant). Legend: AFM = atomic force microscopy; CMG = CellMask™ Green; G-CTL = G-NVs procedural control; G-NVs = ginger-derived nanovesicles; (F-)NTA = (fluorescence) nanoparticle tracking analysis; TEM = transmission electron microscopy.

TEM of G-NVs showed that most of the imaged structures had morphology and size typically attributable to small vesicles, well distinguished from the appearance of the G-CTL, which displayed only a handful of amorphous entities, aggregates, along with a few G-NV-like bodies (Figure 2C). AFM quantitative morphometry [46] also supported these findings (Figure 2D). Although both samples showed abundant globular particles in the AFM images, the vast majority of those found in the G-CTL sample displayed morpho-mechanical characteristics incompatible with intact EVs [41,42]. Conversely, most of the particles contained in G-NVs clustered around a plot region described as characteristic for EVs [42]. In particular, in the G-NV sample, particles above 50 nm diameter (*i.e.*, the size that corresponds to the cut-off of the TFF device used for G-NV enrichment) showed an average CA of 88.0° ± 14.3°, falling within the expected range for EVs (*i.e.*, average CAs of 80° to 120°). In the same sample, we also observed a cluster of smaller objects showing a much higher dispersion of CAs and extending to higher mechanical stiffnesses (*i.e.*, CA > 130°), which were not previously associated with EVs [42], thus suggesting the residual presence of non-vesicular co-isolates. As expected, the G-CTL sample only exhibited the latter type of particles, confirming the selective enrichment of particles smaller than 50 nm and incompatible with the nanomechanical characteristics of intact EVs. AFM morphometry was also employed to assess the relative contribution of different particle diameter ranges to the total particle volume, thus further emphasizing the difference between G-NV and G-CTL. This analysis clearly showed the presence of a substantial fraction of particles larger than 100 nm in the G-NV sample, accounting for more than half of the total particle volume. In contrast, the G-CTL contained only the particles smaller than 100 nm and at much lower prevalence (Figure S1C).

Data collected from the G-NV and G-CTL samples were also validated by fluorescence NTA (F-NTA), which was performed to verify the presence of lipid membranous nanoparticles in our G-NV sample. G-NV particles were labelled with CellMask™ Green (CMG) Plasma Membrane Stain, a lipophilic dye that emits bright and uniform green fluorescence upon insertion into the outer leaflet of lipid membranes. An optimized staining protocol was followed as previously described [39] and SEC was used to purify stained vesicles from the excess of unbound dye, to avoid signals due to dye molecules self-aggregation [46,47]. The sensitivity of the F-NTA instrument required for reliable detection and quantification of fluorescent nanoparticle signals makes the analysis more susceptible to background noise derived from impurities, especially unbound dye [39]. As a control, a procedural blank sample, corresponding to the buffer (PBS) supplemented only with CMG, was processed analogously and assessed in parallel. Averaging the data obtained from the F-NTA analysis of two serial sample dilutions in PBS (1:10 and 1:20), approximately 65 ± 8 % of scattering particles was fluorescent, suggesting that a modest fraction of the total G-NV particle population detected by NTA is non-vesicular. This result is consistent with AFM analysis, which showed that 72% of the detected G-NV particles had a nanomechanical behavior typical for intact vesicles. Using both sample dilutions, the proportion of fluorescent particles relative to total scattering particles did not significantly differ and showed consistent size distribution (Figure 2E). This confirmed that fluorescent labels were stably associated with intact vesicles. Scattering and fluorescently labelled G-NV particles showed overlapping size distributions, indicating stable incorporation of the dye into G-NV vesicle membranes without altering particle size or aggregation state (Figure 2F). Analysis of the blank showed negligible particle count in scatter mode, while revealing a detectable particle population in fluorescence mode, exhibiting a large hydrodynamic diameter (Figure S1D). On one hand, this proved that SEC was effective in removing most of the unbound CMG. On the other hand, it also highlighted that a fraction of the free dye potentially formed self-aggregates or micelle-like structures, which was expected in a neutral and hydrophilic environment as PBS. The wider size distribution of the detected fluorescent particles (Figure S1E) can be potentially attributed to stronger brightness resulting from the clustering of multiple dye molecules, which may lead to size overestimation. F-NTA of the G-CTL yielded particle counts that were comparable to those observed in the PBS (Figure S1D). Nonetheless, size estimation did not follow the same trend, showing not significant differences between scattering and fluorescent particles (Figure S1F). This may confirm the presence of a small subset of residual lipid particles that may sequester CMG, thereby also limiting its self-aggregation or the formation of large micelle-like structures. In contrast to scatter detection, the sensitivity of which depends on particle size and refractive index, fluorescence detection mainly relies on the brightness, stability, and intensity of the fluorophore in use. Therefore, particles that are too small or weak in terms of refractive index to be revealed in scatter mode may be detected in fluorescence mode if conjugated to a proper fluorophore [38]. This explains a higher number of fluorescent particles relative to scattering particles in CMG-labelled G-CTL sample, which may indicate the detection of lipid particles emitting scattering signals falling below the resolution of the instrument that become visible only upon staining.

In alignment with these observations, the G-NV sample exhibited a higher particle number-to-protein ratio, compared with the G-CTL, which is indicative of greater sample purity from protein contaminants (Figure 2G) [47]. As EVs are fundamentally characterized by a presence of a phospholipid bilayer, the protein-to-lipid ratio has been introduced in the EV field as an additional metric that could be used to evaluate the degree of contamination of EV samples from soluble proteins [47,48]. We specifically focused on analyzing the ratio between protein content and phospholipid content, which was observed to be comparable between G-NV and G-CTL preparations, confirming the presence of small phospholipid particles also in the G-CTL, but primarily without typical EV connotation (Figure 2C-D). Furthermore, as *Bt* preferentially utilizes sugars as primary carbon source, we were also interested in quantifying total carbohydrate content in both samples, which highlighted a higher carbohydrate enrichment per protein content in the vesicular preparation (Figure 2I). Overall, these data proved that our manufacturing protocol was efficient in enriching our G-NV sample with particles having typical nanovesicular features and a suitable degree of purity.

### 3.2. IMC-based assessment of *Bt*’s growth under treatment with G-NVs and G-CTL

To evaluate the effect of G-NVs on *Bt*‘s growth, untreated bacteria were analyzed in parallel with cultures treated with three doses of G-NVs: 20,000 particles/cell (dose 1); 10,000 particles/cell (dose 2); 5,000 particles/cell (dose 3) (Table 1). To distinguish the effects specifically attributable to the G-NV preparation from those caused by the ginger matrix separated from it, bacteria were also challenged with the G-CTL at doses normalized to equivalent protein amount of respective G-NV sample dose. Total protein content was selected as normalization factor between vesicles and their respective procedural controls as proteins constitute predominant component of both samples, while particles were observed to be highly enriched in the G-NV preparation. Indeed, the NTA-estimated number of detected particles in G-CTL dose 1, dose 2, and dose 3 corresponded to 60, 30, and 15 particles per cell, respectively. Bacterial growth was monitored via IMC for up to 24 hours (Table 1).

**Table 1.** *Bt* treatment conditions with three doses of G-NVs and G-CTL quantified by NTA. The summary table shows the treatment conditions of *Bt* 1E+7 CFU mL with three doses of G-NVs and G-CTL in terms of NTA-estimated particle concentration – expressed as number of particles per milliliter of treatment reaction, particle size – expressed as mode (nm) and median (nm), and number of particles per treated *Bt* cell. Data are presented as average of four biological batches of G-NVs and G-CTL. Legend: *Bt* = *Bacteroides thetaiotaomicron*; G-CTL = G-NVs procedural control; G-NVs = ginger-derived nanovesicles.

| Sample Name | Dose Number | Particle Concentration | Particle Mode | Particle Median | Inoculum | Particle Number/Treated Cell |
| --- | --- | --- | --- | --- | --- | --- |
| G-NVs | 1 | 2.0E+11/mL | 98.9 ± 7.6 (nm) | 97.5 ± 6.0 (nm) | 1E+7 CFU/mL | 20000 |
|  | 2 | 1.0E+11/mL |  |  |  | 10000 |
|  | 3 | 5.0E+10/mL |  |  |  | 5000 |
| G-CTL | 1 | 6.0E+08/mL | 91.0 ± 7.6 (nm) | 90.1 ± 6.4 (nm) |  | 60 |
|  | 2 | 3.0E+08/mL |  |  |  | 30 |
|  | 3 | 1.5E+08/mL |  |  |  | 15 |

Figure 3A-B illustrates the total heat and heat flow curves of untreated *Bt* and was used as a reference. A parallel analysis of non-inoculated ginger samples confirmed the absence of microbial contamination (Figure S2A-B). The total heat produced by *Bt* (*Q_tot_* = 1.81 ± 0.09 J mL^-1^) was not affected by G-NV supplementation. In contrast, addition of the G-CTL induced a dose-dependent increase in *Q_tot_* (*i.e.*, dose 1: +69%; dose 2: +33%; dose 3: +18%), reaching statistical significance at the two highest doses (Figure 4A). A similar trend was observed for the heat released during the exponential phase (*Q_exp_*), although no statistical difference was detected between bacteria treated with G-NVs dose 3 and its corresponding G-CTL (Figure 4B). IMC data were complemented by biomass measurements obtained via OD_600_ (Figure 4C). The specific growth rate of *Bt* (*µ_max_* = 0.56 ± 0.01 h^-1^) increased significantly in cultures supplemented with either G-NVs or G-CTL, except for the G-CTL dose 3. The increase of *µ_max_* was more pronounced for cultures exposed to G-NVs (*i.e.*, dose 1: +54%; dose 2: +50%; dose 3: +39%), whereas G-CTL treatments exhibited a clearer dose-dependent pattern (*i.e.*, dose 1: +29%; dose 2: +22%) (Figure 4D). A higher maximum metabolic activity level (*P_max_*) was exhibited by all treated cultures (*i.e.*, G-NVs dose 1: +34%; G-NVs dose 2: +32%; G-NVs dose 3: +26%; G-CTL dose 1: +80%; G-CTL dose 2: +47%; CTL dose 3: +34%), compared with control bacteria (*P_max_* = 141 ± 8.91 µW mL^-1^). A statistically significant difference between G-NV samples and their respective G-CTL was observed only at the highest dose, with a stronger response induced by the G-CTL (*i.e.*, +34% with G-NVs dose 1 vs. +80% with G-CTL dose 1) (Figure 4E). The time at which *P_max_* occurred (*T_Pmax_* = 10.8 ± 0.4 h in control bacteria) was significantly shorter with all doses of both G-NVs and G-CTL (*i.e.*, G-NVs dose 1: 7.2 ± 1.2 h; G-NVs dose 2: 7.1 ± 1.3 h; G-NVs dose 3: 7.3 ± 1.4 h; G-CTL dose 1: 8.3 ± 1.6 h; G-CTL dose 2: 8.3 ± 1.6 h; CTL dose 3: 8.0 ± 1.1 h), indicating accelerated growth. This was in line with the thermogram profiles shown in Figure 3. Although no significant difference was found between G-NV- and G-CTL-treated cultures, G-NVs tended to promote slightly faster bacterial growth (Figure 4F).

**Figure 3:**
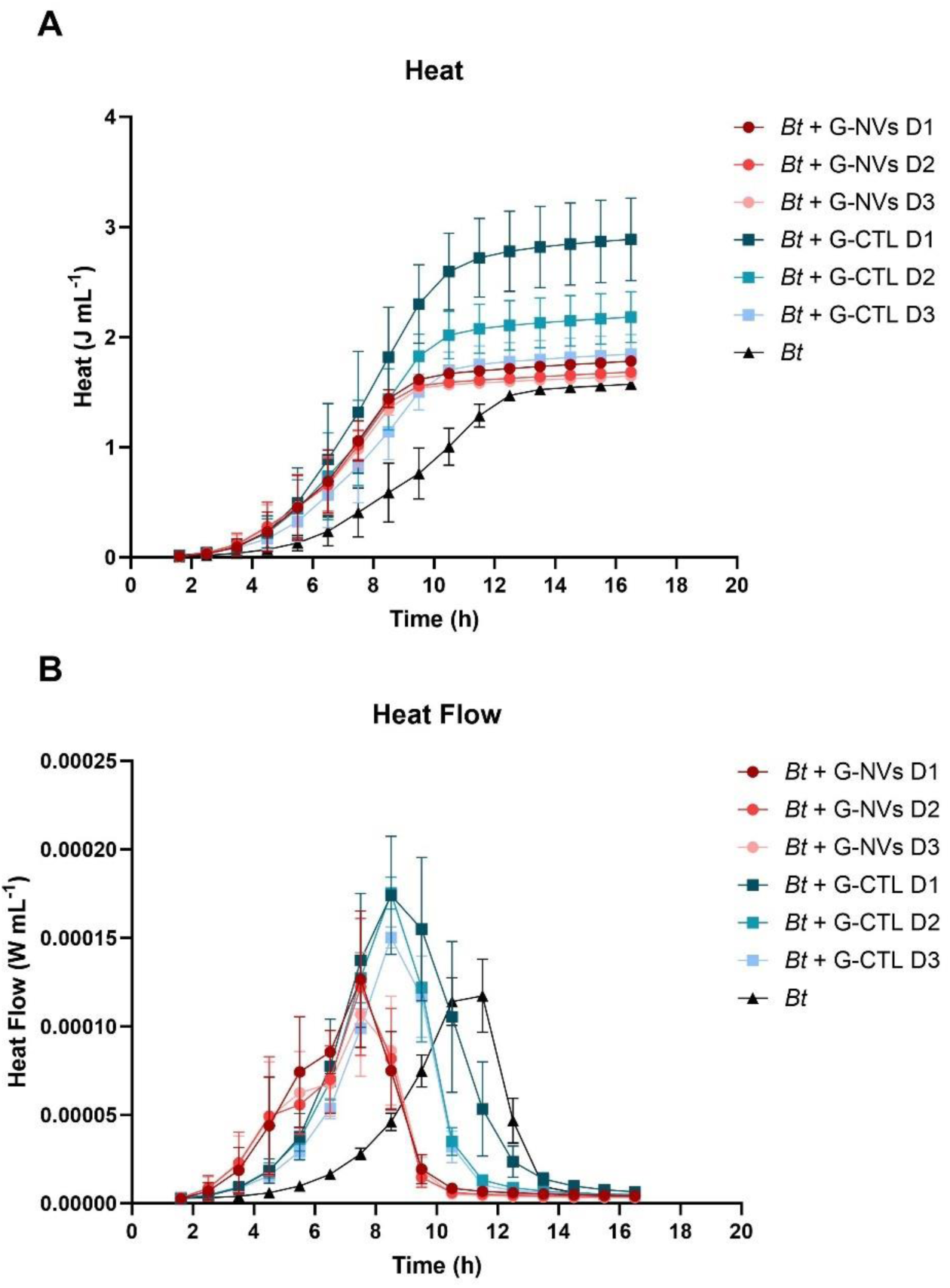
IMC-estimated thermograms of *Bt* grown with or without G-NVs and G-CTL. The line graphs show (A) heat and (B) heat flow curves of *Bt* grown at 37°C for 24h in growth media containing or not containing either G-NVs or G-CTL at dose 1, dose 2 or dose 3. Data are reported as mean ± SEM of four biological batches. Legend: *Bt* = *Bacteroides thetaiotaomicron*; D = dose; G-CTL = G-NVs procedural control; G-NVs = ginger-derived nanovesicles; IMC = isothermal microcalorimetry.

**Figure 4:**
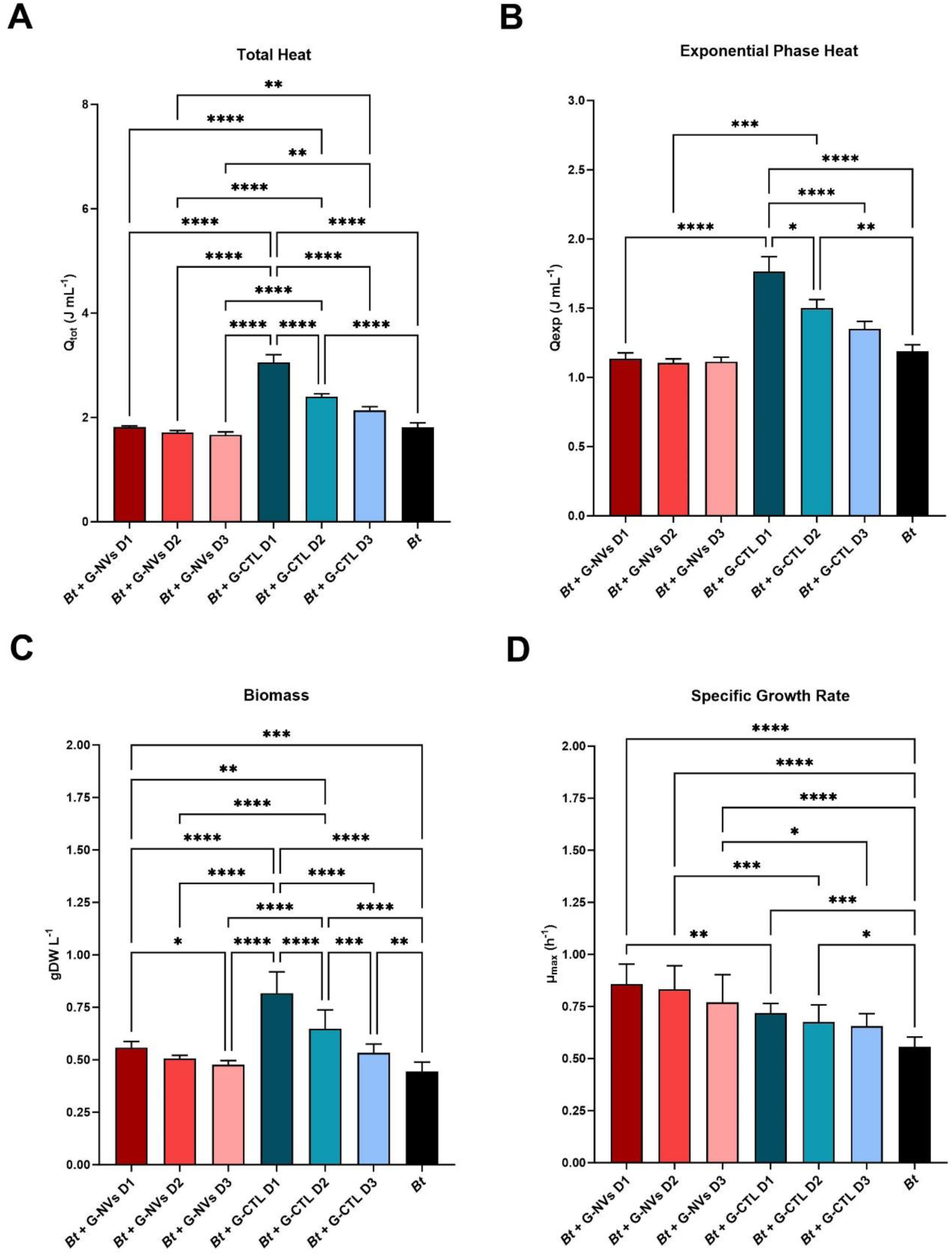
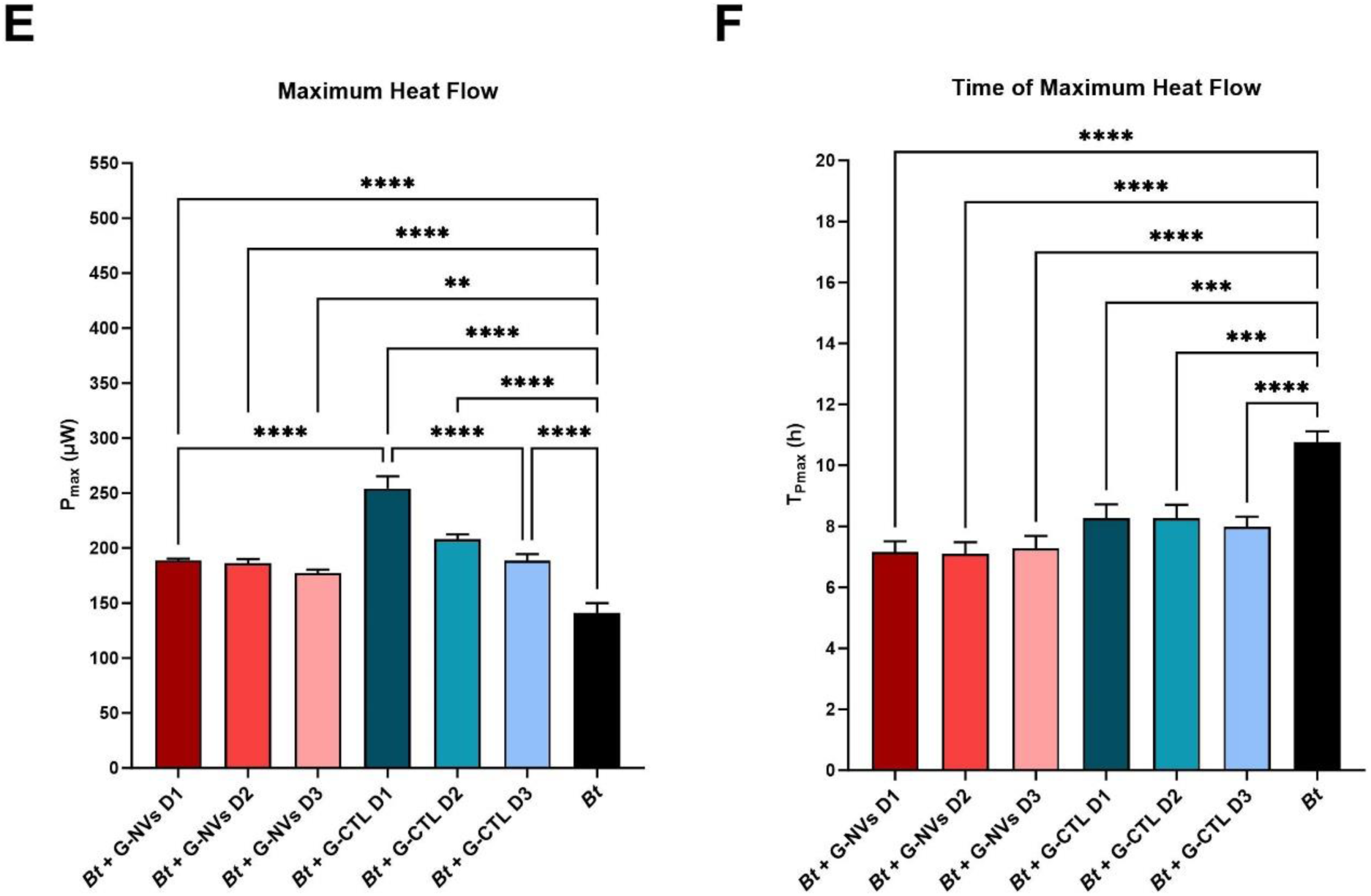
IMC-derived growth characteristics and OD_600_ of *Bt* grown with or without G-NVs and G-CTL. The bar graphs show (A) the cumulative heat produced during the whole cultivation period (*Q_tot_*), expressed as Joule (J) per milliliter, (B) the cumulative heat produced during the exponential phase (*Q_exp_*), expressed as Joule (J) per milliliter, (C) biomass, expressed as grams of dry weight (gDW) per liter, (D) the maximum specific growth rate (*µ_max_*), expressed per (h), (E) the maximum power output at the highest peak of the heat flow curve (*P_max_*), expressed as microwatt (µW), and (F) the time at which the *P_max_* occurs (*T_Pmax_*), expressed as time (h), of *Bt* grown at 37°C for 24h in growth media containing or not containing either G-NVs or G-CTL at dose 1, dose 2 or dose 3. Data are reported as mean ± SEM of four biological batches. Statistical analysis was performed using ordinary one-way ANOVA with follow up test for multiple comparisons (not reported p-value: not significant; significant p-value: < 0.05 [*], <0.01 [**], <0.001 [***], <0.0001 [****]). Comparisons between *Bt* + G-NVs doses and *Bt* + G-CTL doses were restricted to matching doses. Legend: *Bt* = *Bacteroides thetaiotaomicron*; D = dose; G-CTL = G-NVs procedural control; G-NVs = ginger-derived nanovesicles; IMC = isothermal microcalorimetry; OD_600_ = optical density at 600 nm wavelength.

### 3.3. Assessment of *Bt*’s extracellular metabolites released in response to treatment with G-NVs and G-CTL

To evaluate the potential metabolic alterations induced in *Bt* by treatment with either G-NVs or G-CTL, extracellular fermentation substrates and products were quantified in both unconditioned and conditioned growth media by HPLC or UPLC-UV. A specific focus was given to sugars [*i.e.*, Glc, Fru, Man, GlcNAc, Rha], organic acids (*i.e.*, acetate, butyrate, propionate, malate, ethanol, formate, fumarate, glucuronic acid, isobutyrate, isovalerate, lactate, succinate), and FAAs (*i.e.*, Ala, Arg, Asn, Asp, Cys, Glu, Gly, His, Ile, Leu, Lys, Met, Phe, Pro, Ser, Thr, Trp, Tyr, Val), the classes of metabolites that serve as key indicators of microbial metabolism. To facilitate data acquisition and interpretation, the assessment of the samples containing G-NVs and G-CTL was restricted to the highest doses, as these induced the most pronounced and measurable biological effects.

#### 3.3.1. Sugars

Among tested sugars, only Glc and Fru could be reliably quantified. During preparation, growth media were supplemented with Glc to provide a primary carbon source for promoting *Bt*’s growth and metabolism (Section 2.3.1). Therefore, prior to bacterial inoculation and IMC-based cultivation, all media, either containing or not G-NVs and G-CTL, showed detectable levels of Glc.

While the control media and the media including G-NVs had comparable Glc concentrations consistent with the originally added Glc, the media formulated with the G-CTL displayed significantly higher Glc amount (Figure 5 and S3), indicating the contribution to Glc supplementation by the G-CTL content. Similarly, although Fru was not supplemented to the original media, it could be measured in the media containing the G-CTL (Figure 5B and S3B). At the end of the IMC experiment, complete sugar consumption upon fermentation was confirmed in all inoculated samples, while the sugar level in non-inoculated samples remained unaltered with respect to the original media composition (Figure 5 and S3).

**Figure 5:**
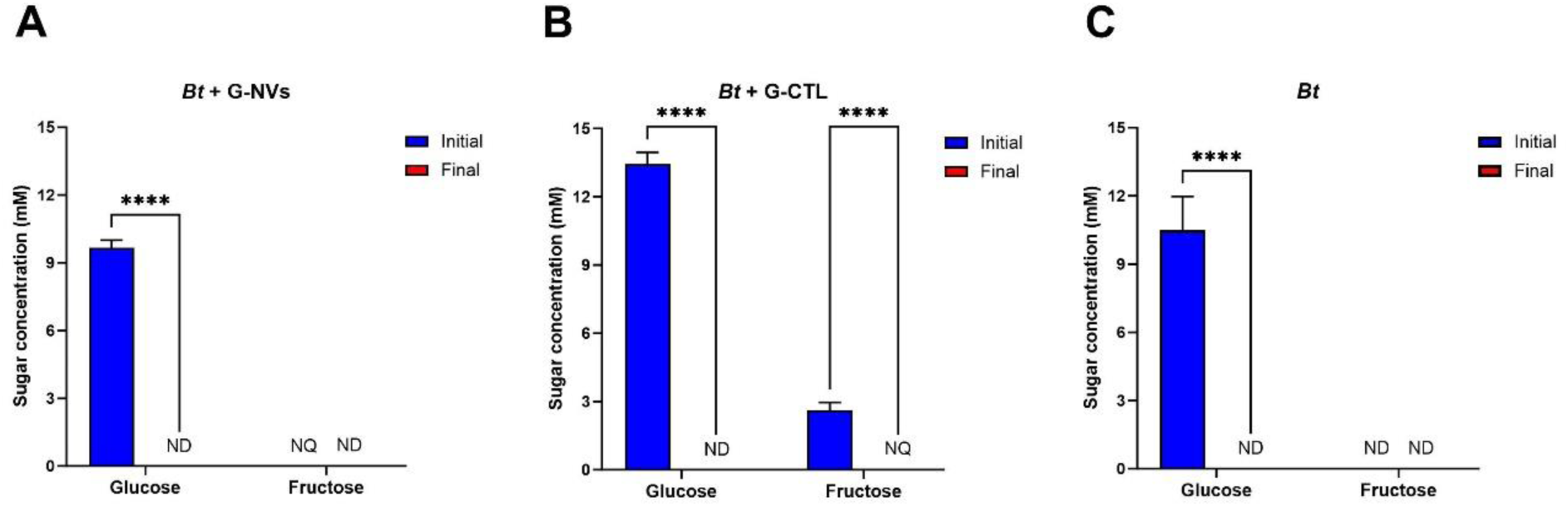
Initial and final free sugar concentrations in *Bt*’s growth media with or without G-NVs and G-CTL. The bar graphs show initial and final sugar concentrations (mM) in *Bt*’s growth media supplemented with (A) G-NVs (dose 1), (B) G-CTL (dose 1), and (C) PBS. Initial concentration data are reported as mean ± SEM of one technical replicate across four independent experiments, each performed using a distinct biological batch of ginger sample. Final concentration data are reported as mean ± SEM of two technical replicates across four independent experiments, each performed using a distinct biological batch of ginger sample. Statistical analysis was performed using ordinary two-way ANOVA with follow up test for multiple comparisons (not reported p-value: not significant; significant p-value: <0.0001 [****]). Legend: *Bt* = *Bacteroides thetaiotaomicron*; G-CTL = G-NVs procedural control; G-NVs = ginger-derived nanovesicles; ND = not detected; NQ = not quantified.

#### 3.3.2. Organic acids and ethanol

Organic acids and ethanol represent primary bacterial fermentation products. Therefore, they were expected to be measured only in the conditioned media collected at the end of bacterial growth. This was confirmed for all tested organic compounds, except for ethanol, which was surprisingly detected in the starting media supplemented with both G-NVs and G-CTL before inoculation with bacteria (Figure 6A-B and S4). To exclude the association of this readout to any artefact, a confirmative GC-MS analysis was performed on replicates of two samples collected at the end of the IMC-based cultivation period, validating our observations by HPLC: i) blank media (not inoculated) enriched with G-NVs; ii) conditioned media, depleted of grown bacteria, that were originally supplemented with the G-CTL (Figure S5). Fermentation of *Bt* induced measurable secretion of ethanol, acetic, formic, propionic, and succinic acids. Among them, acetic, succinic, and formic acids revealed the highest levels. A significantly greater production and release of acetic, succinic, and propionic acids was observed in samples supplemented only with G-CTL, while enrichment of growth media with G-NVs did not provide any added value in boosting bacterial metabolic activity (Figure 6B and S4). The end concentrations of ethanol, normalized with initial concentrations, and formic acid did not reveal any significant differences between control and treated bacteria (Figure S6).

**Figure 6:**
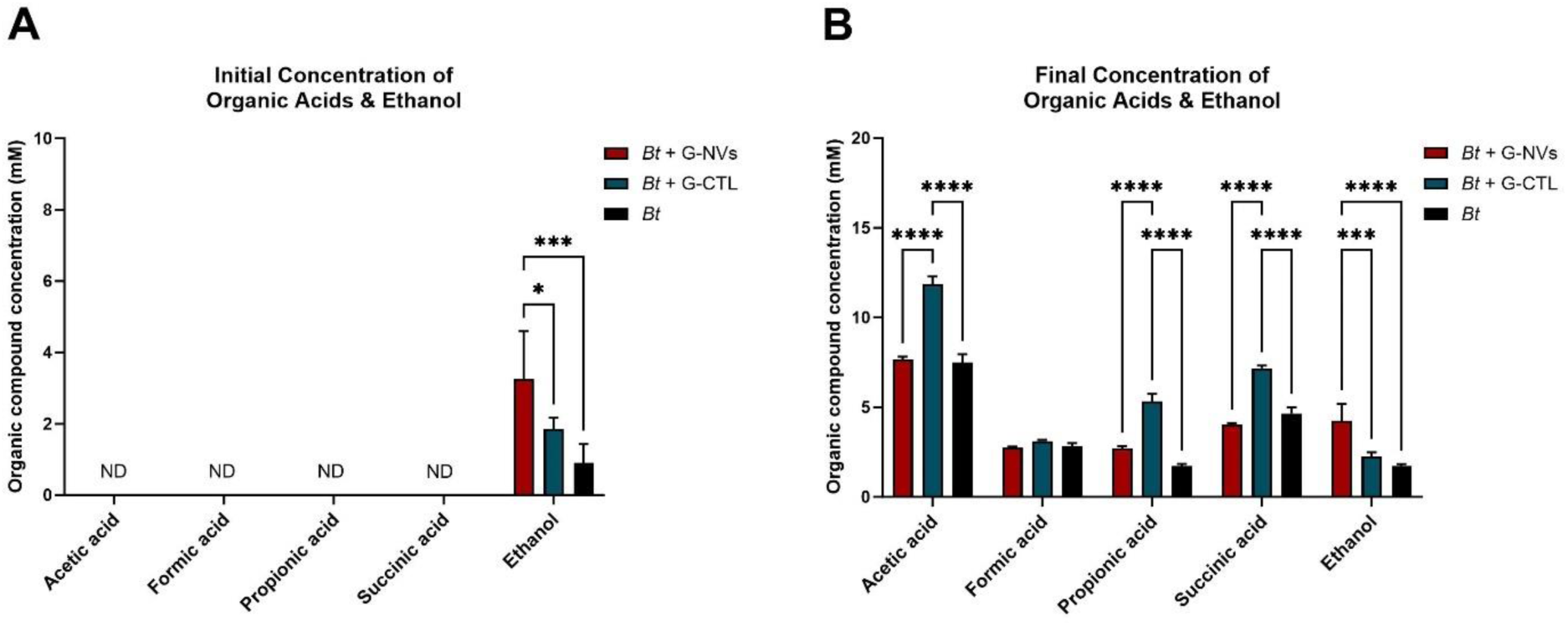
Initial and final concentrations of organic acids and ethanol in *Bt*’s growth media with or without G-NVs and G-CTL. The bar graphs show (A) initial and (B) final concentrations (mM) of organic acids and ethanol in (A) non-inoculated media and (B) inoculated media. Data in A are reported as mean ± SEM of one technical replicate across four independent experiments, each performed using a distinct biological batch of ginger sample. Data in B are reported as mean ± SEM of two technical replicates across four independent experiments, each performed using a distinct biological batch of ginger sample. Statistical analysis was performed using ordinary two-way ANOVA with follow up test for multiple comparisons (not reported p-value: not significant; significant p-value: < 0.05 [*], <0.001 [***], <0.0001 [****]). Legend: *Bt* = *Bacteroides thetaiotaomicron*; G-CTL = G-NVs procedural control; G-NVs = ginger-derived nanovesicles; ND = not detected.

#### 3.3.3. FAAs

Initial concentrations of all tested FAAs were shown to be comparable across all growth media before inoculation (Figure 7A and S7). Interestingly, a significantly lower level of NH_4_+ was observed in the media containing G-CTL, compared with the control media and the media supplemented with G-NVs (Figure 7A). Although *Bt* is known to be both a saccharolytic and a proteolytic microbial species [12], no changes in FAA concentrations were observed in any sample at the end of the cultivation period (Figure 7B and S7). Unexpectedly, a significant reduction in NH_4_+ was also highlighted in the non-inoculated medium containing G-NVs post IMC experiment (Figure S7D).

**Figure 7:**
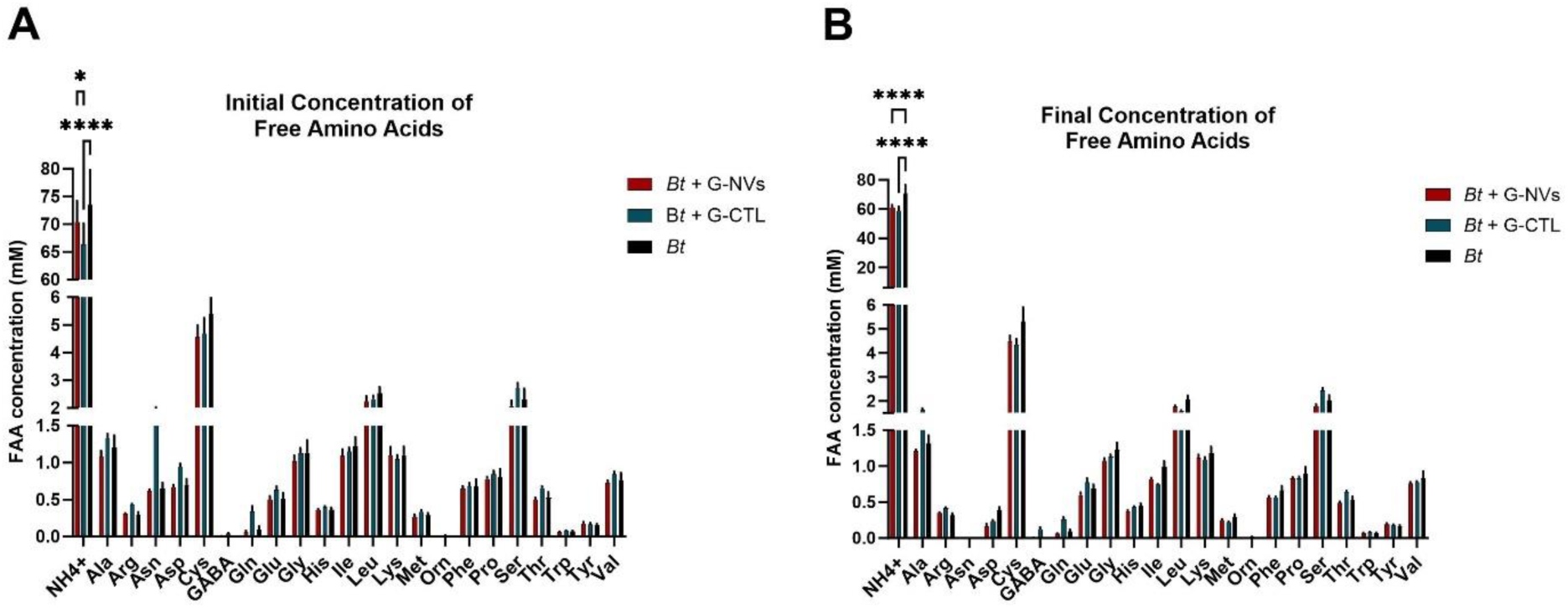
Initial and final FAA concentrations in *Bt*’s growth media with or without G-NVs and G-CTL. The bar graphs show (A) initial and (B) final concentrations (mM) of FAAs in (A) non-inoculated media and (B) inoculated media. Data in A are reported as mean ± SEM of one technical replicate across four independent experiments, each performed using a distinct biological batch of ginger sample. Data in B are reported as mean ± SEM of two technical replicates across four independent experiments, each performed using a distinct biological batch of ginger sample. Statistical analysis was performed using ordinary two-way ANOVA with follow up test for multiple comparisons (not reported p-value: not significant; significant p-value: < 0.05 [*], <0.0001 [****]). Legend: *Bt* = *Bacteroides thetaiotaomicron*; G-CTL = G-NVs procedural control; G-NVs = ginger-derived nanovesicles.

## 4. Discussion

In the present study, we explored the growth and metabolic response of *Bt*, an obligate anaerobe of the HGM with encouraging exploitable potential as NGP [18,19], to the exposure with small nanovesicles derived from ginger rhizome (*Zingiber officinale Roscoe*), hereby indicated as G-NVs. In preclinical models, G-NVs have been shown to modulate the microbial composition of the HGM, actively inducing anti-inflammatory and hepatoprotective effects [49,50]. However, to our knowledge, this work showcases the first application of IMC to probe the impact of G-NVs, and EVs in general, on bacterial growth dynamics, enabling kinetic measurement of biomass production and real-time monitoring of metabolic activity. Such methodology may provide an appealing tool for a currently unmet need in studying EV roles and applications in microbiology. As recommended by the “Minimal Information for Studies of EVs” (MISEV) 2018 and 2023 guidelines [46,47], we have assessed G-NVs against a procedural control sample, which we named G-CTL, to distinguish the biological activity attributable to the 50-200 nm sized particles enriched in our G-NV preparation, from the residual functionality of the ginger extract depleted of the G-NV and containing constituents smaller than 50 nm.

Ginger samples were prepared using a combination of state-of-the-art methodologies for EV production, including differential centrifugation, dead-end filtration, and TFF. In the final processing step, both G-NVs and G-CTL were recovered as the retentate and permeate, respectively, from a TFF device with a 50 nm pore-size membrane, followed by subsequent sterilization by passage through a 0.22 µm pore-sized dead-end filter unit. For their physicochemical characterization, we adopted complementary biophysical and biochemical approaches, integrating single-particle, imaging, and bulk biochemical analyses. NTA, selected as a widely used and well-established high-throughput method for single-particle analysis and quantification [46], showed that the G-NV sample had a substantial concentration of particles within the expected size range for nanovesicles. NTA is measuring the hydrodynamic diameter of particles in solution based on their Brownian motion, thus potentially leading to particle size overestimation [46]. This may account for the apparent, though significantly reduced particle concentration (nearly 800-fold lower), detected in the G-CTL preparation, exhibiting a size range partially overlapping with the particle size distribution estimated in the G-NV sample. To validate either the presence or the absence of lipid membranous particles in our G-NV and G-CTL preparations, respectively, both were stained with CMG, a lipophilic dye emitting green fluorescence upon insertion into lipid membranes, and analyzed by F-NTA. Additionally, low-throughput imaging technologies, specifically TEM and AFM, were adopted to complement F-NTA analysis by enabling the detection of particles falling below the detection limit of the NTA instrument, and providing insights into the morpho-nanomechanical properties of particles. With respect to the G-NV sample, these methods confirmed the enrichment of the G-NV sample in particles displaying characteristic vesicular morphology and membrane stiffness within the anticipated size distribution of 50-200 nm. It also revealed the presence of a residual subset of particles, either vesicular or showing amorphous appearance and increased stiffness, likely corresponding to co-isolated contaminants, such as protein aggregates and polysaccharide fragments. Likewise, these analyses corroborated the NTA measurements of the G-CTL, highlighting the selective accumulation of small particles, especially those falling below the 50 nm diameter threshold, most of which exhibited higher stiffness than vesicles, as well as amorphous shape. Protein and phospholipid quantification were in line with the observed enrichment of nanovesicles in the G-NV sample, which exhibited higher concentrations of phospholipidic particles purified from soluble proteins, compared with the G-CTL. We observed minimal retention of particles smaller than the ultrafilter cut-off, as well as passage of particles bigger than the filter cut-off into the permeate, a phenomenon largely documented in ultrafiltration processes, likely due to diverse physicochemical and operational factors. The correlation between the nominal cut-off attributed to ultrafiltration systems and their actual retention efficiency is indeed governed not only by physical size exclusion, but also by the hydrodynamic behavior of particles and the hydrodynamic drag force, which regulate particle-particle and particle-surface interactions, ultimately affecting flow dynamics, filtering membrane fouling, and particle separation efficiency [51–53]. Furthermore, it is known that particles characterized by softer membranes and greater flexibility, such as EVs, may deform and squeeze through smaller pores into the permeate [42,54]. Overall, our isolation protocol yielded G-NV samples with a significant enrichment in EV-like nanoparticles with a suitable degree of purity and exhibiting distinct physicochemical features, compared with the G-CTL preparation.

Our experimental setting comprised the dose-dependent response of *Bt* to ginger samples, assessed by treating bacterial cells with G-NV and G-CTL preparations at corresponding doses normalized to equivalent total protein amount. Along with EV particle number, this parameter is among the most common normalization factors used for EV functional assessment *in vitro* and *in vivo* [55]. Protein content can be reliably quantified in both samples, reflecting the molecular sample makeup. Proteins, along with lipids, are the predominant components of EVs, forming their structure and carrying out functions [56,57].

*Bt*’s treatment with G-NVs, at concentrations of up to 20,000 particles per cell, did not affect the heat released during either the whole *c*ultivation period (*Q_tot_*) or solely the exponential growth phase (*Q_exp_*), suggesting no alteration of the total biomass output. This was validated by traditional OD_600_ measurement and further supported by metabolite profiling, which showed no changes in fermentation end-products, such as SCFAs and FAAs. These findings concur with evidence from a recent study, reporting no impact of G-NVs on the individual growth of *Bacteroides fragilis*, used as a reference microbial strain of the Bacteroidaceae family [50]. The G-CTL was analyzed in parallel to assess *Bt*’s response to ginger non-vesicular components. G-CTL elicited a specific boost in fermentation yield, as reflected by the induced dose-dependent increase in heat outputs, both *Q_tot_* and *Q_exp_* and OD_600_ readings. The greater and dose-dependent secretion of acetic, succinic, and propionic acids promoted by the G-CTL confirmed its stimulatory effect. Although microbial fermentation under G-NV conditioning resulted in unaltered cellular expansion and release of extracellular metabolites, we observed that both G-NVs and G-CTL enhanced *Bt*’s specific growth rate (*µ_max_*), with G-NVs exerting a more substantial effect, raising the maximum metabolic activity output (*P_max_*) and inducing earlier occurrence of the *P_max_* (*T_Pmax_*). This indicates that both ginger samples triggered an early metabolic response, but resulting in different metabolic output, which was steady with G-NVs while increased by the G-CTL. As *Bt* is preferentially involved in the decomposition of indigestible, high molecular weight (HMW) carbohydrates (*e.g.*, dietary fibers and host mucus-derived glycans) [10–12], the greater carbohydrate-to-protein ratio quantified in the G-CTL, with respect to G-NVs, likely contributed to the observed functional discrepancy between the two samples. In support of this, no consumption of FAAs was observed under any tested condition, confirming *Bt*’s prioritization of carbohydrates as a primary carbon source for meeting metabolic demand.

The higher specific growth rate promoted by G-NVs in comparison with the G-CTL, is potentially attributable to differences in multiple mechanisms, including cargo delivery and activation of cell signaling pathways. The G-NV-mediated co-transport of carbohydrates with other active biomolecules, such as proteins, lipids, and nucleic acids, as well as the potential induction of signaling cascades by ligand-receptor interactions on bacterial cell surface without internalization, may exert rapid and synergistic effects, leading to the simultaneous activation of complementary metabolic and regulatory pathways, thus accelerating overall bacterial cellular response [58,59]. In *Bacteroides* spp., the internalization of soluble polysaccharides (PSs) is primarily mediated by the starch utilization system (Sus), which is localized in the bacterial outer membrane. This apparatus functions in coordination with cell surface and periplasmic glycosidases, which digest HMW-PS, and the inner membrane transport system, which promotes the final entry of oligosaccharides and monosaccharides into the cytoplasm [60,61]. Due to the complex and diversified nature of this mechanism, the rate of carbohydrate processing and uptake mediated by the Sus may be highly affected by various factors. These include the PS MW, structure, local concentration, and cellular energic state, as transport of carbohydrates against their electrochemical gradient is energy intensive. The process may be also influenced by prior exposure history, as cells previously exposed to carbohydrates maintain the Sus ready for new exposure, thus shortening its induction latency [62].

Different EVs are reported to induce rapid functional and metabolic changes through mechanisms that do not require full cargo internalization by the recipient or interacting cell, such as binding to surface receptors or through the delivery of cargo that directly interacts with the cell surface or the surrounding extracellular environment. However, EV-mediated metabolic changes also and frequently rely on vesicle uptake via membrane fusion or endocytosis, processes that enables the internalizing cell to access the EV’s internal cargo [59,63]. While EV uptake can occur rapidly (within minutes), the rate can also vary substantially depending on the interacting EV types and recipient cells. The rapid and transient “kiss-and-run” contacts between EVs and target cell membranes have been described in human cell systems. These events can mediate either cell signaling or fusion accompanied by cargo release, both potentially influencing downstream cellular responses [64]. In gram-negative bacteria, EV uptake has been shown to be relatively slow, occurring after 2 hours of treatment. This rate is slower than the bacteria division rate and is influenced by factors such as vesicle composition, size, bacterial growth phase, and the specific mechanisms used for uptake, which, in the case of PDVs, remain uncertain [65,66]. Although both membrane fusion and endocytosis are energy-dependent processes, EVs, including PDVs, may increase the rate of cargo internalization and utilization bypassing the lag phase required by the Sus to sense and pre-digest the substrates [66,67].

Beyond glycosylated biomolecules, vesicles may be already loaded with soluble carbohydrates or contain active glycosidases that catalyze localized substrate hydrolysis in specific microenvironments – either in proximity to cells, within vesicles, or at contact zones between vesicles and substrate [68–70]. Furthermore, by enabling the direct delivery of enclosed and concentrated cargo by fusogenic or endocytotic pathways, they can ensure controlled intracellular release and prevent random diffusion to the surrounding environment [71–73]. It was interesting to observe that the addition of either G-NVs or G-CTL to the non-inoculated incubation media induced a significant and comparable reduction of NH_4+_, which was faster with the G-CTL. This may denote that both ginger samples contain metabolic enzymes incorporating NH_4+_ into organic compounds (*e.g.*, glutamine synthetase), or that they sequester NH_4+_, potentially due to dynamic electrostatic interactions with macromolecules, such as nucleic acids, making it undetectable in free form [74–76]. Nevertheless, solid evidence supporting these hypotheses is still lacking.

The unexpected detection of ethanol in the same set of samples, not linked to microbial contamination, as demonstrated by both IMC and OD_600_ measurements, could be the result of partial contamination by microbial biomolecules, such active enzymes, and substrates, involved in microbial fermentation. Because of the known symbiotic relationship of ginger with other microorganisms [77], these could have been co-isolated with ginger components upon sample preparation, although the contamination with viable bacteria was excluded. Alternatively, ethanol production could have been directly promoted by ginger in response to the activation of an endogenous plant ethanolic fermentation pathway, which typically involves two key enzyme, pyruvate decarboxylase and alcohol dehydrogenase, that may have been collected in our ginger samples in association with vesicles or in soluble form [78]. Overall, these assumptions warrant further investigation for confirmation.

## 5. Conclusions

IMC demonstrated that both G-NVs and G-CTL affect *Bt*’s growth and metabolic response, although to varying extents and mediating different cargo delivery mechanisms. G-NVs showed to provide faster and more efficient metabolic stimulation, without increasing total metabolic output. Conversely, the soluble biocomponents enriched in G-CTL caused a sustained, dose-dependent metabolic effect, due to the higher cargo availability and potentially promoting cargo uptake through alternative pathways. The obtained data also highlighted the competitive advantage of IMC over conventional methods for assessing changes in bacterial growth dynamics and kinetics following incubation with EVs, including PDVs, thereby offering a novel and strategic approach to advance research and development focusing on (plant)EV/NV-microbe interactions.

## 6. Perspectives and Future Directions

The use of IMC in this study revealed a critical gap in the current EV research landscape, requiring elucidation of the mechanisms of action of PDVs with cells and, in general, EVs with bacteria. The analysis of the dynamic metabolic effects induced by EV-microbe interactions is often overlooked. From the parallel physicochemical and functional characterization of G-NVs and G-CTL within the selected microbiological model for bioactivity testing, we identified novel insights into the structural and functional differentiation between vesicular and non-vesicular components derived from ginger. The observed increase in *Bt*’s specific growth rate in response to treatment with both G-NVs and G-CTL, in association with a greater fermentation yield induced by the latter, advances our understanding of how ginger extracts exert different metabolic effects, depending on the relative contribution of bioactive constituents, as well as whether they are delivered in soluble form or within vesicles. Our approach and findings can also integrate present methods for fine-tuning *Bt* and overall gut microbial activity, thereby advancing current fermentation design and process optimization strategies, while also fostering the development of nutraceutical products – not only NGPs, but also prebiotics (*i.e.*, substances providing health benefits after utilization by microorganisms) and synbiotics (*i.e.*, combined probiotics and prebiotics) – and microbiome-targeted therapies [20,79–82]. For instance, our ginger preparations, or other samples with similar properties, may be exploited to prime and accelerate bacterial metabolism. Moreover, PDVs sharing characteristics with G-NVs may serve as efficient delivery systems of active nutrients or for therapeutics targeting of the HGM. On the other hand, systematic profiling of substrates promoting greater microbial metabolic output, such as our G-CTL, may support prebiotic design.

To support mechanistic understanding, our future studies will be dedicated to conducting high-throughput multi-omics profiling of tested ginger samples and *Bt*’s secretome, while extending biomass and metabolic assessment to IMC intermediate time-points. To deepen *Bt*’s metabolic response efficiency and dynamics, we also aim at exploring how the secretion and composition of *Bt*’s own EVs, conventionally known as Outer Membrane Vesicles (OMVs) [83], are modulated by G-NVs and G-CTL, along with elucidating triggered uptake mechanisms. Finally, we plan to expand IMC research to additional PDVs and NGP models tested either in monocultures or within synthetic, multi-strain consortia [16,84], thereby enhancing our understanding not only on PDV-mediated plant-microbe interactions, but also on microbial cross-feeding mechanisms and community stability.

## 6. Glossary of terms

Ala: Alanine
NH_4+_: Ammonium
Arg: Arginine
Asn: Asparagine
Asp: Aspartic Acid
AFM: Atomic Force Microscopy
Bt: Bacteroides thetaiotaomicron
BCA: Bicinchoninic Acid
Cys: Cysteine
CMG: CellMask™ Green Plasma Membrane Stain
CFU: Colony-forming Units
CA: Contact Angle
*Q_exp_*: Exponential (Phase) Heat
EV: Extracellular Vesicle
FI: Fluorescence Intensity
F-NTA: Fluorescence Nanoparticle Tracking Analysis
FAA: Free Amino Acid
Fru: Fructose
Glu: Glutamic Acid
gDW: Grams of Dry Weight
G-NVs: Ginger-derived Nanovesicles
G-CTL: Ginger-derived Nanovesicles Control
Glc: Glucose
Gly: Glycine
(H)MW: (High) Molecular Weight
HPLC: High-Performance Liquid Chromatography
His: Histidine
HGM: Human Gut Microbiota
Ile: Isoleucine
IMC: Isothermal Microcalorimetry
Leu: Leucine
Lys: Lysine
Man: Mannose
*P_max_*: Maximum Power Output
*µ_max_*: Maximum Specific Growth rate
Met: Methionine
MISEV: Minimal information for studies of extracellular vesicles
GlcNAc: N-acetyl-D-glucosamine
NTA: Nanoparticle Tracking Analysis
NGP: Next-generation Probiotic
ND: Not Detected
NQ: Not Quantified
OD: Optical Density
OMV: Outer Membrane Vesicle
Phe: Phenylalanine
PBS: Phosphate-Buffered Saline
PDV: Plant-derived Vesicle
PS: Polysaccharide
Pro: Proline
Rha: Rhamnose
RT: Room Temperature
Ser: Serine
SCFA: Short Chain Fatty Acid
SEM: Standard Error of Mean
Sus: Starch Utilization System
TFF: Tangential Flow Filtration
Thr: Threonine
*T_Pmax_*: Time of *P_max_*
*Q_tot_*: Total Heat
TEM: Transmission Electron Microscopy
Trp: Tryptophan
Tyr: Tyrosine
UPLC-UV: Ultra Performance Liquid Chromatography-UV
Val: Valine

## 7. CRediT authorship contribution statement

**Francesca Loria**: Conceptualization; Methodology; Investigation; Validation; Formal Analysis; Data Curation; Visualization; Writing–Original Draft Preparation. **Anna Kattel**: Methodology; Investigation; Validation; Visualization; Writing–Review & Editing. **Marco Brucale**: Resources; Investigation; Writing–Review & Editing. **Francesco Valle**: Resources; Investigation. **Paolo Bergese**: Writing–Review & Editing. **Eeva-Gerda Kobrin**: Resources; Investigation; Validation; Data Curation; Writing–Review & Editing. **Antonio Chiesi**: Funding acquisition; Resources. **Paolo Guazzi**: Writing–Review & Editing. **Sirin Korulu**. Writing–Review & Editing; Supervision. **Irina Stulova**. Resources; Formal Analysis; Data Curation; Writing–Review & Editing. **Nataša Zarovni**: Conceptualization; Methodology; Resources; Writing–Review & Editing; Supervision. **Raivo Vilu**: Funding acquisition; Conceptualization; Methodology; Resources; Writing–Review & Editing; Supervision.

## 8. Acknowledgements

This work was funded by the following projects: i) “Fermentation-based Legume Protein Extraction Technology” (KAVATECH) project (Project No: RE.5.04.24-0655); ii) “Harnessing the Microbial Potential of Fermented Foods for Healthy and Sustainable Food Systems” (DOMINO) project (Project No: 101060218); iii) the European Commission-funded H2020-FETOPEN-2016-2017 "The Extracellular Vesicle Foundry" (evFOUNDRY) project (Grant Agreement ID: 801367); iv) the European Commission-funded H2020-FETPROACT-2019-2020 "Biogenic Organotropic Wetsuits" (BOW) project (Grant Agreement ID: 952183). We acknowledge the services of the University of Helsinki: HiPREP Core in FIMM Technology Centre supported by HiLIFE and Biocenter Finland for performing electron microscopy work and Electron Microscopy Unit of the Institute of Biotechnology for providing the facilities. We also thank TFTAK AS for providing opportunity to analyze the samples. We are particularly grateful to Scientists Marina Junusova, Merli Špitsmeister, and Aaro Videvik for kindly performing, respectively, sugar, organic acids, and ethanol analysis, FAA analysis, and a confirmative GC-MS analysis.

## 9. Conflict of interest statement

The company HansaBioMed Life Sciences Ltd, current employer of Francesca Loria, Sirin Korulu, Paolo Guazzi, and Antonio Chiesi is the manufacturer of some of the products used in this study: TFF-EVs-Small device, Phospholipid Quantification Assay Kit, black and transparent PS microplate wells with no binding capacity. The authors declare that they have no conflict of interest.

