## Supplementary Information for "Investigating the effect of ginger-derived nanovesicles on the growth and metabolic activity of *Bacteroides thetaiotaomicron*: an isothermal microcalorimetric study"

### 1. Supplementary Information

#### 1.1. Supplementary Figures

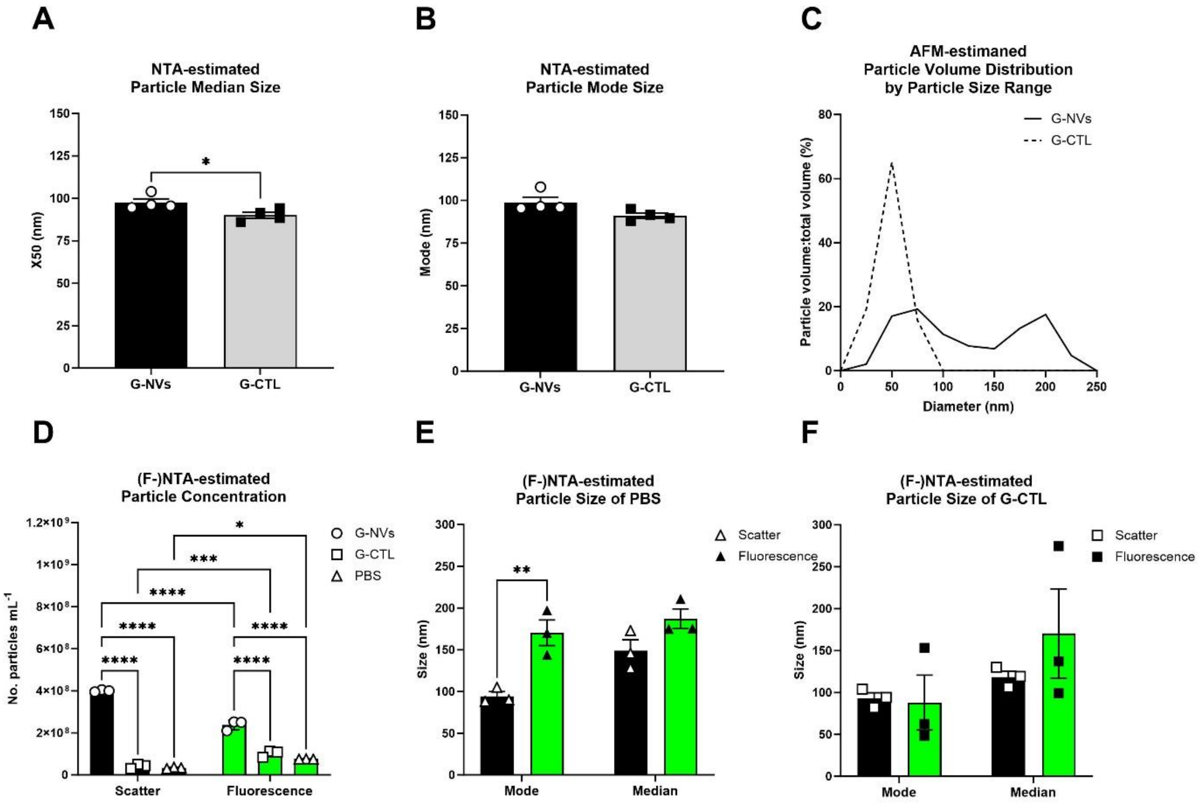


**Figure S1: Physicochemical assessment of G-NVs, G-CTL, and procedural controls**. The bar graphs show: A) NTA-estimated particle median size (nm) of four biological batches of G-NVs and G-CTL; B) mode size (nm) of four biological batches of G-NVs and G-CTL; D) (F-)NTA-estimated particle concentrations of three batches of CMG-labelled G-NVs, G-CTL, and PBS – expressed as particle number per milliliter – measured with 1:10 dilution in PBS; E) (F-)NTA-estimated particle mode size (nm) and median size (nm) of three biological batches of CMG-labelled G-CTL measured in scattering and fluorescence mode with 1:10 dilution in PBS; F) (F-)NTA-estimated particle mode size (nm) and median size (nm) of three replicates of CMG-labelled PBS measured in scattering and fluorescence mode with 1:10 dilution in PBS. The histograms in C depict the percentage fractions of total particle volume divided by particle diameter range of one representative batch of G-NVs and G-CTL, estimated by liquid AFM. Data in A and B are reported as mean ± SEM of four biological batches of G-NVs and G-CTL. In A and B, statistical analysis was performed using ordinary unpaired T test (not reported p-value: not significant; significant p-value: < 0.05 [*], <0.0001 [****]). In D, statistical analysis was performed between matching scatter and fluorescence values, as well as between different samples within either scatter or fluorescence measurements, using ordinary two-way ANOVA with follow up test for multiple comparisons (significant p-value: <0.01 [**], <0.001 [***], <0.0001 [****]). In E and F, statistical analysis was performed between matching scatter and fluorescence values using ordinary two-way ANOVA with follow up test for multiple comparisons (significant p-value: <0.01 [**], <0.001 [***]). Legend: AFM = atomic force microscopy; CMG = CellMask™ Green; G-CTL = G-NVs procedural control; G-NVs = ginger-derived nanovesicles; (F-)NTA = (fluorescence) nanoparticle tracking analysis.

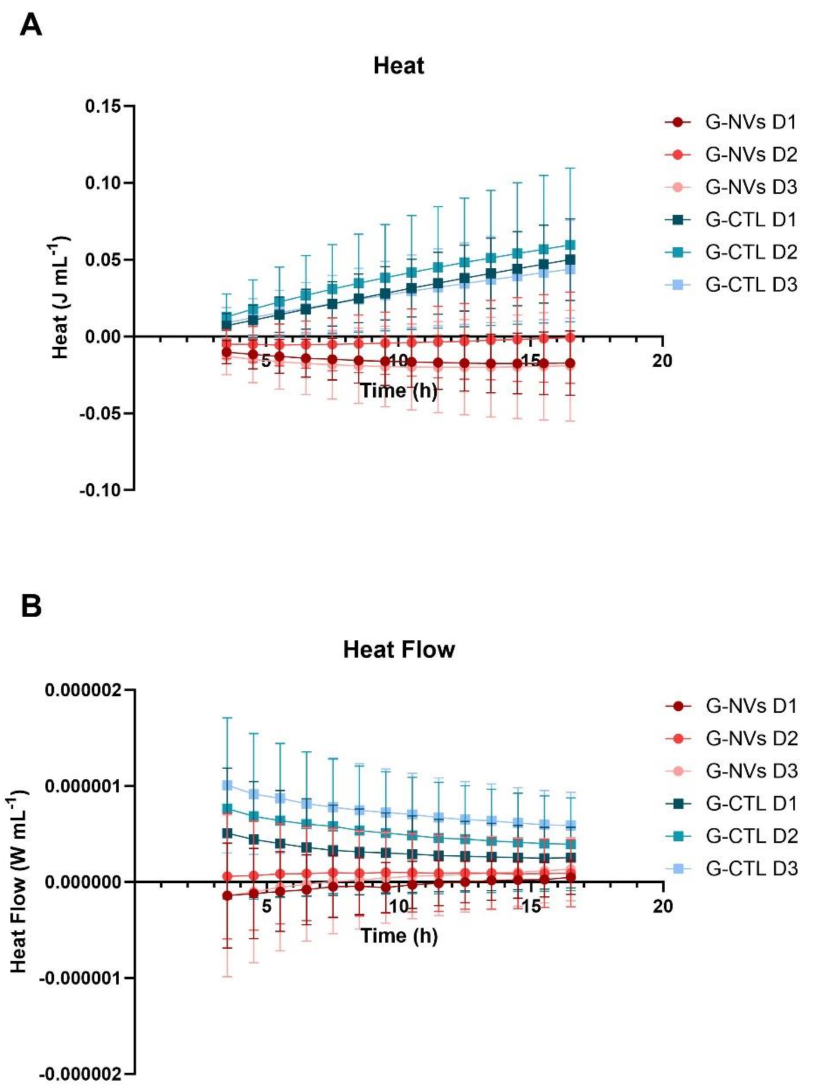


**Figure S2: IMC-estimated thermograms of non-inoculated *Bt*'s growth media with G-NVs or G-CTL.** The line graphs show (A) heat and (B) heat flow curves of non-inoculated *Bt*'s growth media supplemented with G-NVs or G-CTL, incubated at 37ºC for 24h. Data are reported as mean ± SEM of four biological batches. Legend: D = dose; G-CTL = G-NVs procedural control; G-NVs = ginger-derived nanovesicles; IMC = isothermal microcalorimetry.

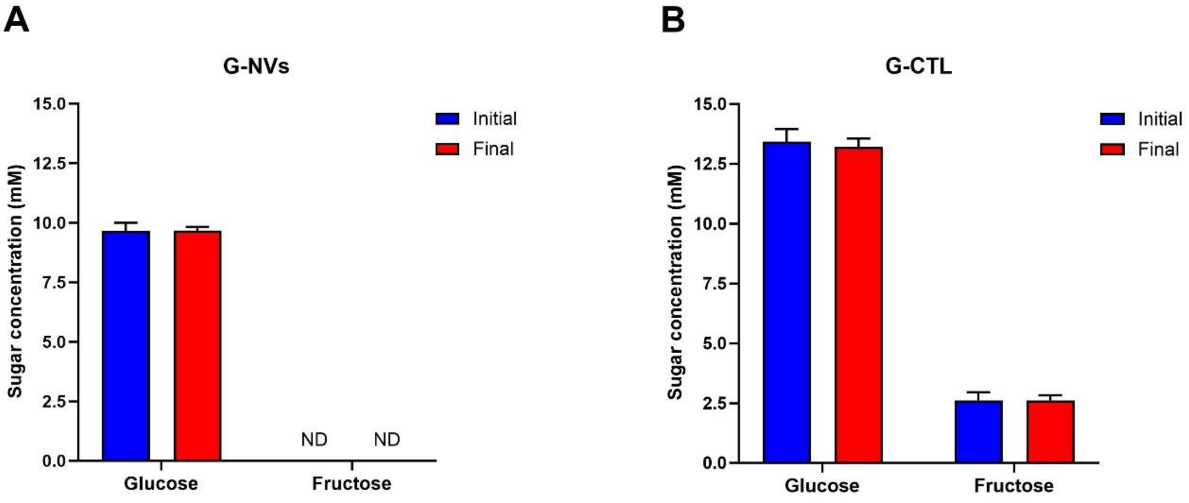


**Figure S3: Initial and final free sugar concentrations in non-inoculated *Bt*'s growth media with G-NVs or G-CTL.** The bar graphs show initial and final sugar concentrations (mM) in non-inoculated *Bt*'s growth media supplemented with (A) G-NVs (dose 1) or (B) G-CTL (dose 1). Initial concentration data are reported as mean ± SEM of one technical replicate across four independent experiments, each performed using a distinct biological batch of ginger sample. Final concentration data are reported as mean ± SEM of two technical replicates across four independent experiments, each performed using a distinct biological batch of ginger sample. Statistical analysis was performed using ordinary two-way ANOVA with follow up test for multiple comparisons (not reported p-value: not significant). Legend: G-NVs = ginger-derived nanovesicles; G-CTL = G-NVs procedural control; ND = not detected.

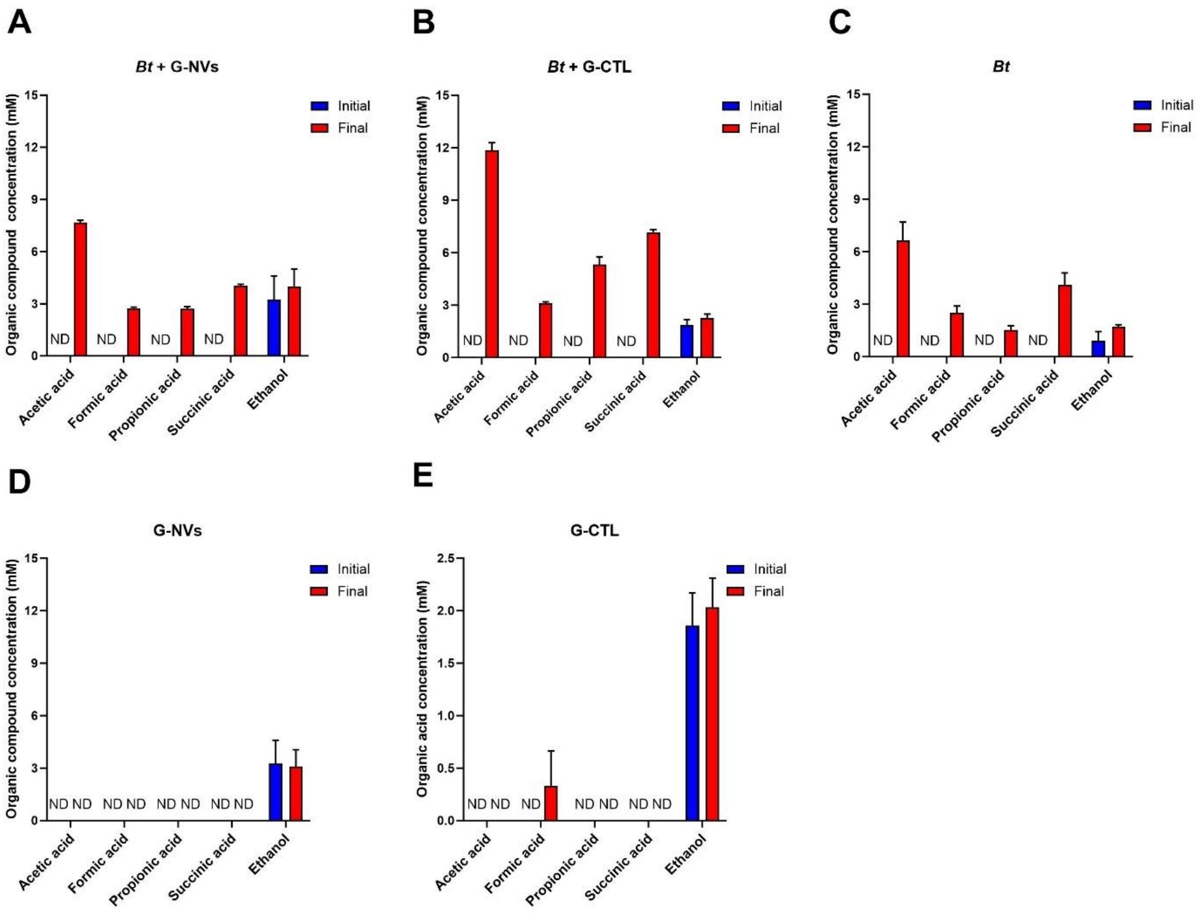


**Figure S4: Initial and final concentrations of organic acids and ethanol in inoculated and non-inoculated *Bt*'s growth media with or without G-NVs and G-CTL.** The bar graphs show initial and final concentrations (mM) of organic acids and ethanol in inoculated (A-C) and non-inoculated (D-E) *Bt*'s growth media supplemented with G-NVs (dose 1) or (B) G-CTL (dose 1). Initial concentration data are reported as mean ± SEM of one technical replicate across four independent experiments, each performed using a distinct biological batch of ginger sample. Final concentration data are reported as mean ± SEM of two technical replicates across four independent experiments, each performed using a distinct biological batch of ginger sample. Statistical analysis was performed using ordinary two-way ANOVA with follow up test for multiple comparisons (not reported p-value: not significant). Legend: *Bt* = *Bacteroides thetaiotaomicron*; G-NVs = ginger-derived nanovesicles; G-CTL = G-NVs procedural control; ND = not detected.

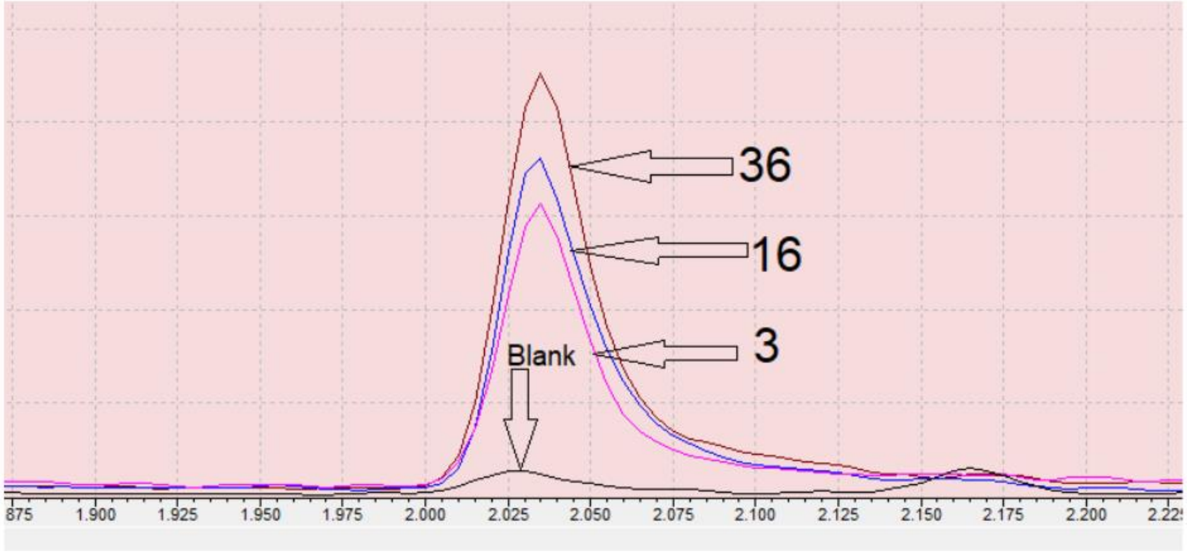


**Figure S5. Control GC-MS analysis.** The GC-MS chromatograms show detectable levels of ethanol in non-inoculated *Bt*'s growth media with G-NVs (dose 1) (sample no. 36) and in fermented *Bt*'s growth media with G-CTL (dose 1) (sample no. 3 and 16). Ethanol was identified with NIST17 library, which gave hit of 98% accuracy. Legend: D = dose; GC-MS = gas chromatography-mass spectrometry; G-CTL = G-NVs procedural control; G-NVs = ginger-derived nanovesicles; IMC = isothermal microcalorimetry.

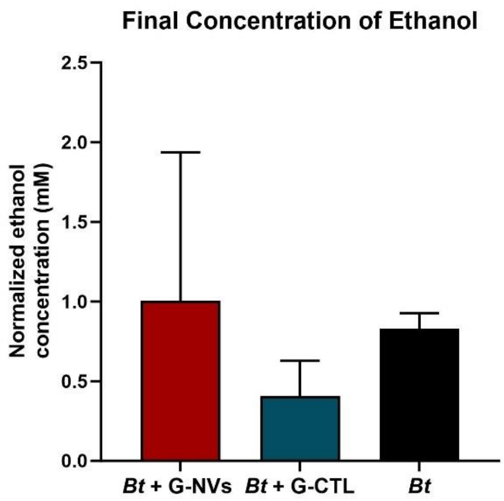


**Figure S6: Normalized ethanol concentrations in fermented *Bt*'s growth media with or without G-NVs and G-CTL.** The bar graph depicts final ethanol concentrations (mM) in *Bt*'s growth media with or without G-NVs (dose 1) and G-CTL (dose 1) after subtraction of initial concentrations. Data are reported as mean ± SEM of two technical replicates relative to four independent experiments, each performed using a distinct biological batch of ginger sample. Statistical analysis was performed using ordinary one-way ANOVA with follow up test for multiple comparisons (not reported p-value: not significant). Legend: *Bt* = *Bacteroides thetaiotaomicron*; G-CTL = G-NVs procedural control; G-NVs = ginger-derived nanovesicles.

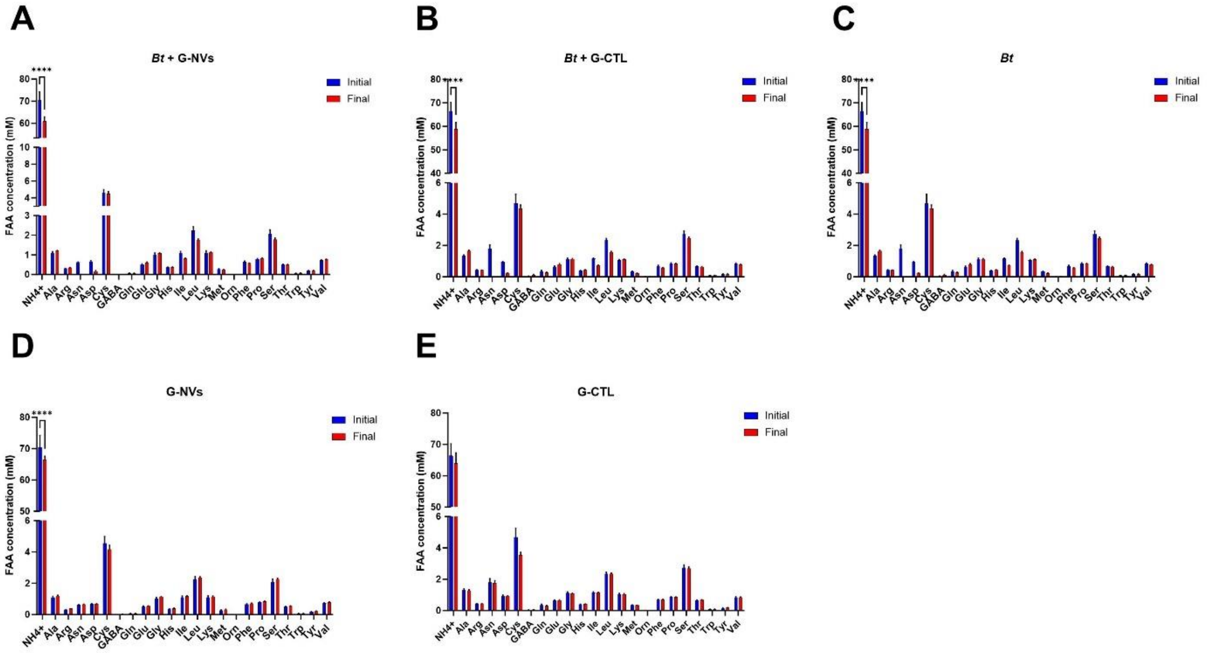


**Figure S7: Initial and final FAA concentrations in *Bt*'s growth media with or without G-NVs and G-CTL.** The bar graphs show initial and final concentrations (mM) of FAAs in (A-C) inoculated and (D-E) non-inoculated *Bt*'s growth media supplemented with G-NVs (dose 1), (B) G-CTL (dose 1), or PBS. Initial concentration data are reported as mean ± SEM of one technical replicate across four independent experiments, each performed using a distinct biological batch of ginger sample. Final concentration data are reported as mean ± SEM of two technical replicates across four independent experiments, each performed using a distinct biological batch of ginger sample. Statistical analysis was performed using ordinary two-way ANOVA with follow up test for multiple comparisons (not reported p-value: not significant; significant p-value: <0.0001 [****]). Legend: *Bt* = *Bacteroides thetaiotaomicron*; G-CTL = G-NVs procedural control; G-NVs = ginger-derived nanovesicles.

#### 1.2. Supplementary Materials and Methods

##### 1.2.1. Gas chromatography–mass spectrometry (GC–MS)

Volatile compounds were extracted using solid-phase microextraction (SPME). A 100 µL aliquot of the sample was transferred into a 10 mL glass vial. A 1 cm SPME fiber coated with 30/50 µm DVB/Car/PDMS (Stableflex) was exposed to the headspace for 40 minutes at 40°C to allow adsorption of volatiles. The collected compounds were then thermally desorbed in the injection port of a gas chromatograph for 5 minutes. Ethanol identification was performed using a Shimadzu GC-2030 system coupled with a 8050NX Triple Quadrupole mass spectrometer (Shimadzu, Kyoto, Japan). Separation was achieved on a ZB5-MS column (30 m × 0.25 mm i.d., 1.0 µm film thickness; Phenomenex, Torrance, CA, USA) with helium as the carrier gas at a linear velocity of 35 cm/s. The oven temperature program started at 40°C, increased at 5°C/min to 190°C, then ramped at 25°C/min to 280°C, with a final hold of 4 minutes, resulting in a total run time of 36 minutes. Mass spectra were acquired using electron ionization at 70 eV, scanning a mass-to-charge (m/z) range of 35–250. Each sample was analyzed in duplicate. Ethanol was identified in a non-targeted manner using GC-MS solution software (Shimadzu) and retention indices (RI), which were calculated based on the retention times of adjacent n-alkanes. Compound identification was confirmed by comparing experimental RI values and spectra with those in the NIST17 and FFNSC spectral libraries.
